# Cytokinin Mediates Photosynthesis and Senescence Action Through CHLOROPLAST IMPORT APPARATUS 2 (CIA2) and CIA2-like (CIL)

**DOI:** 10.64898/2026.09.30.755754

**Authors:** Risheek R. Khanna, Satyam Singh, Miroslav Berka, Shelby Crowell, Peyton Hass, Alvaro Sanz-Saez, Martin Černý, Aaron M. Rashotte

## Abstract

Progression of leaf senescence is regulated by internal factors like the plant hormone cytokinin (CK) that acts to delay senescence and retain photosynthesis by maintaining chloroplast-functionality. While initial CK-signaling components controlling senescence have been described, downstream targets regulating physiological performance have remained less characterized. Here we present the CHLOROPLAST IMPORT APARATUS 2 (CIA2) and CIA2-like (CIL) transcription factors (TFs) as mediators of CK senescence activity, enabling chloroplast-function. Utilizing loss-of-function mutants, we show CIA2 and CIL negatively regulate senescence progression and alter physiological performance in *Arabidopsis thaliana*. Integrated transcriptomic-metabolomic analysis shows that CIA2 and CIL action is distinct from other known chloroplast-related TFs (like GLKs/GATAs) and can boost antioxidant response over senescence. We find that CIA2 and CIL are essential in relaying CK activity during chloroplast biogenesis, photosynthesis, rubisco function, and metabolite accumulation. In turn, ARR10/12 CK signaling components were needed for CK induction of *CIA2* and *CIL* themselves - a cascade that induced chloroplast- and photosynthesis-associated nuclear genes, to boost chloroplast-function and metabolism. We also show that CK acts to repress senescence through accumulation of the polyamine, spermidine, in a CIA2/CIL dependent manner. This work presents a CIA2/CIL-cascade as a new anterograde signal for CK antisenescence effects, indicating a mechanism through which CK prolongs leaf function. This fundamental understanding of CK action has implications towards engineering plants with elevated photosynthesis performance and delayed senescence.

## Introduction

Senescence represents the final stage of leaf development yet controls critical activities to ensure that emerging organs receive essential recycled nutrients and sugars from this senescing tissue (Lim et al., 2007; Woo et al., 2019; Guo et al., 2021). The process of senescence is regulated by both exogenous cues and endogenous factors such as the plant hormone cytokinin (CK) that delays it (Argueso and Kieber, 2024). CKs are plant hormones known to regulate chloroplast development and leaf senescence (Zwack and Rashotte, 2013; Cortleven and Schmülling, 2015). *trans*-Zeatin (*t*Z) is a naturally occurring CK form that is highly active in eliciting an antisenescence response and improving chlorophyll retention, resulting in greener leaves for longer (Khanna et al., 2025, Hasannin et al., 2026; Thennakoon et al., 2026). Thus, senescence can be simply observed using an established CK bioassay, where CK application delays dark-induced leaf senescence (DIS; Letham, 1967, Johnston et al., 2026). DIS mimics developmental senescence and is often used to examine senescence responses in addition to CK effects (Paluch-Lubawa et al., 2021). Processes occurring during senescence include reactive oxygen species (ROS) production leading to oxidative stress damage, membrane disintegration, photosynthesis machinery breakdown, and loss of chlorophyll pigments (Bresson et al., 2018). During this process, various senescence-associated genes (SAGs) are induced and act as molecular markers of onset and progression of senescence (Breeze et al., 2011).

CK is perceived by binding Histidine Kinase (AHK) receptors that initiate a two-component signaling cascade (TCS), eventually activating two types of Response Regulators (RR), type-B RRBs that function as TFs and type-A RRAs that function as negative regulators of CK-signaling (Argueso and Kieber, 2024). Cytokinin Response Factors (CRFs) are also TF that operate as side branches of CK signaling interacting with RRs (Hallmark and Rashotte, 2019). Previous work on senescence has demonstrated a primary role for three CK signaling components in this process: the receptor AHK3, the RRB ARR2, and CRF6 (Kim et al., 2006; Zwack and Rashotte, 2013). While several studies have focused on characterizing gene/protein regulation of senescence (reviewed in Guo et al., 2021), these studies have not attempted to identify downstream processes and targets of CK signaling TFs. As such, the mechanism of CK action that maintains physiological performance in senescence is not well understood.

The chloroplast is a sub-cellular organelle involved in photosynthesis and metabolism. Communication between the nucleus and chloroplast is essential for biogenesis and maintenance of the plastid (Mahapatra et al., 2024). Most chloroplast-associated proteins are nuclear-encoded and post-translationally imported through a TOC/TIC import complex (Xing et al., 2025). Photosynthesis and chloroplast-function are essential for normal leaf developmental events including senescence. Chloroplast-function is directly linked to leaf senescence as it is clearly involved in photosynthetic processes and ROS generation. As senescence progresses a breakdown of photosystem apparatus and chloroplast thylakoid membranes occurs, while CK acts to reduce this damage (Carillo and Ferrante, 2025; Kábrtová et al., 2026). Additionally, CK is known to regulate chloroplast development and chloroplast-associated genes involved in organelle and pigment biogenesis, and photosynthesis processes (Cortleven and Schmülling, 2015). Chloroplast Import Apparatus 2 (CIA2) is an important TF protein that induces gene expression crucial for import and assembly of chloroplasts, photosystem functioning, and chloroplast-related protein synthesis (Sun et al., 2001, 2009). CIA2-like (CIL) shares high sequence identity with CIA2 and functional redundancy (Yang and Sun, 2020). KO *cia2 cil* double mutants result in reduced chlorophyll pigment, altered chloroplast ultrastructure, and limited protein import, overall being detrimental to chloroplast activity (Li et al., 2021; Gawroński et al., 2021; Yang et al., 2022).

Part of how CK functions to maintain chloroplast-functionality (or anterograde signaling) is through the transcriptional regulation of chloroplast- and photosynthesis-associated nuclear genes (*Cp-* & *PhANG*s) - either directly through RRBs or indirectly through other chloroplast-related TF such as Golden2-like (GLK) and GATA TFs (Cackett et al., 2022). *GLK* genes are founding members of the GARP (Golden2, ARR-B and PSR1) TF family and the *glk1*,*2* double mutant KO results in altered chloroplast ultrastructure and functionality, and a smaller rosette diameter, thus closely resembling *cia2 cil* phenotype (Waters et al., 2009; Li et al., 2021). Related GATA TFs (GATA21/GNC) and CK-responsive GATA1 (GATA22/CGA1) also regulate chloroplast development and greening of plants, again with a *gata21,22* double mutant exhibiting pale phenotypes (Chiang et al., 2012). While these GLKs and GATAs can transcriptionally control chloroplast biogenesis, very few TFs that regulate the *Cp-* & *PhANG*s have been studied in the context of senescence.

Here we show that exogenous CK application enhances chloroplast biogenesis, photosynthesis, and photosynthate reserve accumulation in a CIA2, CIL dependent manner in *Arabidopsis thaliana*. Thus, CIA2 and CIL are mediators in CK improvement of chloroplast-functionality and antisenescence. We then show the induction of *CIA2* and *CIL* by CK is explained through its regulation by TCS components, *RRB*s, particularly *ARR10/12*. Further, using transcriptomics and co-expression analysis we show that *CIA2* and *CIL*, mediate a CK anterograde signal that is crucial for normal senescence progression and distinct from known chloroplast-related factors (*GLK*s/*GATA*s). Finally, using metabolite profiling we show that CK induces accumulation of chloroplast-related metabolites like spermidine, a polyamine that represses leaf senescence. Together, this work presents a CIA2/CIL-cascade that CK utilizes towards maintaining chloroplast-function and retaining physiological performance under senescence.

## Results

### CIA2 and CIL influence plant growth, photosynthesis, and physiology

CK can regulate chloroplast function, however the cascades mediating such activity remain less explored, which we predicted could be controlled through CIA2 and CIL to relay CK activity. Prior to examining CK based effects, we assessed CIA2 and CIL gene function and the extent to which they control growth, photosynthesis, and physiology. Here we analyzed the physiological consequences of loss-of-function in *cia2-1* (hereafter *cia2*)*, cil-5* (hereafter *cil*), and *cia2-1 cil-5* (hereafter *dm*) mutants in Arabidopsis (Fig 1). *cia2* exhibits a pale phenotype with a 24% reduction in chlorophyll content relative to WT (Fig 1A,B), similar to that seen by Li et al., 2021 under different conditions. *cil* resembled WT pigment content, however the *dm* showed a paler phenotype compared to all other genotypes (43% reduction relative to WT) – indicating functional redundancy in CIA2 and CIL.

**Figure 1:**
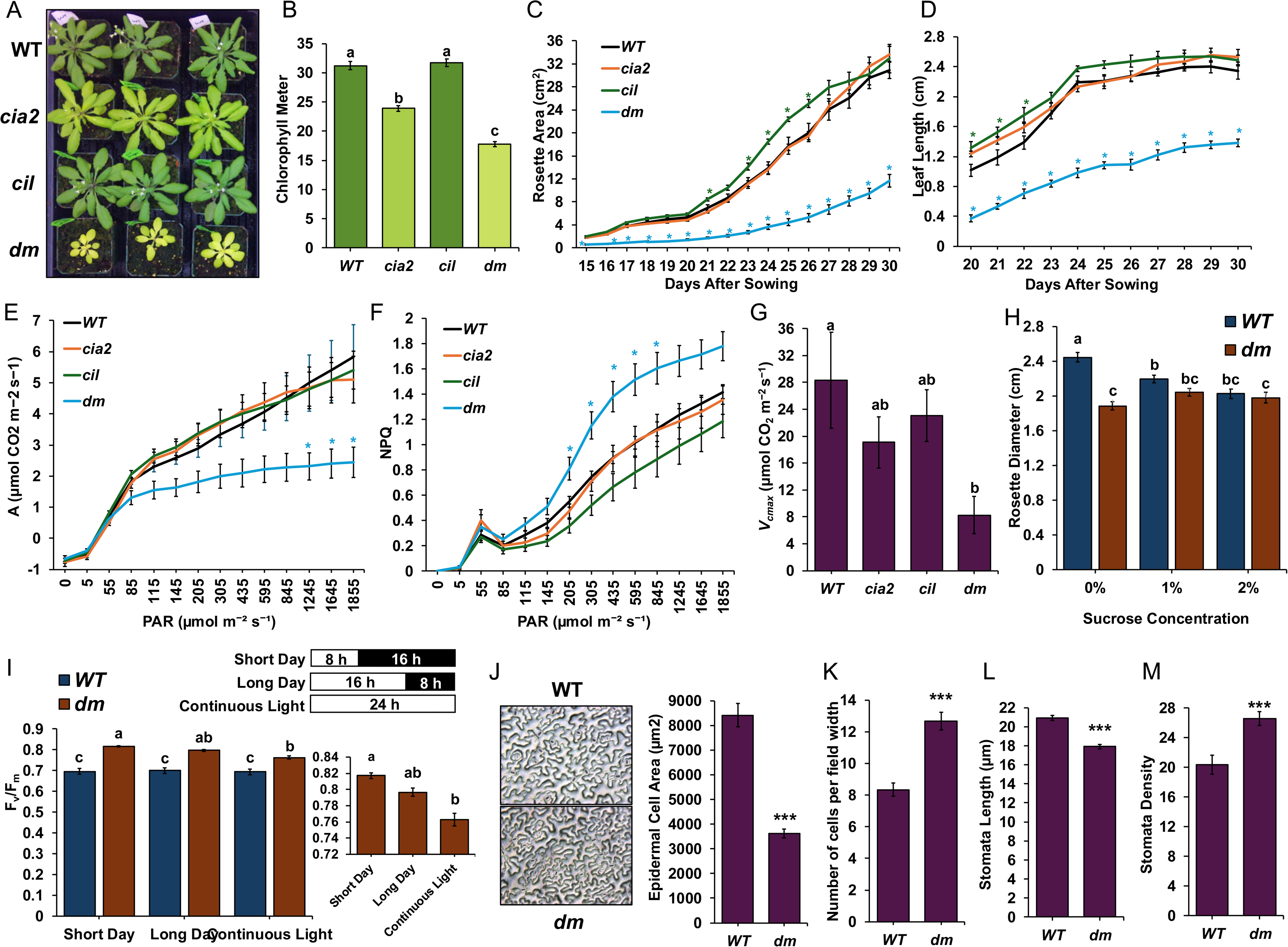
CIA2 and CIL influence plant growth, photosynthesis, and physiology. (A) WT (Col-0), *cia2-1* (CS6522; hereafter *cia2*), *cil-5* (CS2110093; hereafter *cil*) and double mutant (CS2110094; hereafter *dm*) genotypes 30 days after sowing (DAS). (B) Chlorophyll levels measured using chlorophyll meter from leaf 5 at 30 DAS (n = 30). (C-D) Image-based phenotyping measuring rosette area (in cm^2^) or leaf 5 length (in cm; n = 6). (E) Photosynthesis (*A*; µmol CO_2_ m^-2^ s^-1^) analysis or (F) non-photochemical quenching (NPQ) across photosynthetically active radiation (PAR; µmol m^-2^ s^-1^). (G) Maximum rate of rubisco carboxylation (*V_cmax_*; μmol CO_2_ m^−2^ s^−1^). Values normalized to 25 °C before presenting. Leaf 5-8 were utilized for all LI6800-based photosynthesis analysis using at least 6 plants per genotype at 45 DAS. (H) Seedlings 5 DAG were moved to MS media with 0%, 1%, or 2% sucrose for 7 days following which rosette diameter was recorded using ImageJ analysis (n = 24). (I) Photosystem II efficiency (F_v_/F_m_) recorded after 7 days of short day, long day, or a continuous photoperiod, measuring at least 30 seedlings. Mini bar graph represents F_v_/F_m_ levels only for *dm* seedlings. (J-M) Abaxial epidermal peel of leaf 5 at 30 DAS plants (20x objective magnification) with representative pictures shown. Three microscopic fields (leaf tip, mid, and base) were captured and epidermal cell size (in µm^2^) measured using ImageJ analysis (measuring 60 cells per genotype) and number of cells across microscopic field width (measuring 6 epidermal peels per genotype). Stomata length (µm) measured using ImageJ analysis (measuring 90 stomata per genotype) and number of stomata per field (18 fields per genotype). All results are represented as mean ±SE for at least three biological replicates. Statistical tests performed using ANOVA and Tukey’s HSD with different letters noting statistically significant difference (p.adj < 0.05) or asterisk (* p.adj < 0.05). Student’s t test performed for (J, K, L, and M) with asterisk (*** p < 0.005).

We further evaluated multiple growth-related parameters (Fig 1, Supp File 1). Using image-based phenotyping, rosette size and diameter of *dm* exhibit a reduced rate of growth (Fig 1C, Supp Fig 1A). WT, *cia2*, and *cil* each showed a rapid increase in growth starting ∼20-21 DAS, which did not occur in *dm* (Fig 1C). Evaluation of leaf 5 length and width suggested that mutations in the *dm* impairs regular leaf development (Fig 1D, Supp Fig 1B). To further ensure growth trends were specifically due to loss of CIA2 and CIL function, we evaluated an independent set of T-DNA insertion alleles: crossing *cia2-2* with *cil-1* and found the *cia2-2 cil-1* also had a reduced rosette diameter (by 29%; Supp Fig 1C).

To understand genotype-effects related to photosynthesis, we performed photosynthesis light and *AC_i_* curve analysis (Fig 1, Supp File 1). Without light, at 0 photosynthetically-active radiation (PAR; μmol m^-2^ s^-1^), all genotypes exhibited photosynthesis (*A*) levels below 0 μmol CO_2_ m^-2^ sec^-1^, indicating no baseline photosynthesis only respiration (Fig 1E). Subsequently under higher PAR exposure above 1245 μmol m^-2^ s^-1^ PAR, *dm* showed significantly reduced photosynthesis relative to WT (Fig 1E). Using a light-curve based modeling approach (Kaipiainen 2009, Lobo et al., 2013), *I_50_* and *I_comp_*levels were evaluated, however no significant difference was observed across genotypes (Supp Fig 1D,E). *dm* showed higher non-photochemical quenching (NPQ) levels, being statistically different at 205, 305, 435, 595, and 845 μmol m^-2^ s^-1^ PAR (Fig 1F). This suggests a reduced ability of light utilization towards photochemistry in *dm*. However, *cia2* and *cil* did not show significantly altered photosynthesis or NPQ in comparison with WT, indicating a CIA2-CIL redundancy towards maintaining photosynthesis only revealed in *dm* (Fig 1E,F). Using *AC_i_* curves, we examined maximum rubisco carboxylation (*V_cmax_*; μmol CO_2_ m^-2^ s^-1^) and found *dm* to have 71% reduced *V_cmax_* levels (Fig 1G). However, the rate of electron transport (*J_max_*; μmol electrons m^−2^ s^−1^) was not significantly different across genotypes (Supp Fig 1F). Overall, this indicates that the rubisco activity is lower in the *dm*, suggesting consequences in light-independent cycles of photosynthesis. For rate of transpiration (*Emm*; mmol m^-2^ s^-1^), single mutant genotypes resembled WT however *dm* exhibited 38% reduced levels (relative to WT) although not significant difference was observed (Supp Fig 1G).

We hypothesized that reduced *dm* chloroplast-function limits photosynthate production, thus impacting growth. To address this, WT and *dm* were grown on media supplemented with 0-2% sucrose (Fig 1H, Supp Fig 1H). *dm* had a smaller rosette compared to WT at 0% sucrose, however not with 1% or 2% sucrose – suggestive of sucrose growth restoration in *dm* to WT levels (Fig 1H). To uncouple this from potential osmotic effects and the pale phenotype displayed by *dm*, we tested a non-destructive photosystem II (PSII) efficiency parameter – F_v_/F_m_ under 0-2% sucrose across the whole rosette and young leaves. Results showed no influence of sucrose towards PSII efficiency (Supp Fig 1I) or towards the pale phenotype of *dm* (Supp Fig 1H). Beyond sugar, we evaluated the influence of photoperiod on the *dm* phenotype (Fig 1I, Supp Fig 1J). *dm* was pale with 31% reduced growth even under short day conditions (8:16 h light:dark; Supp Fig 1J) suggesting CIA2 and CIL as necessary to retain growth irrespective of photoperiod. While *dm* had higher F_v_/F_m_ levels compared to WT across all photoperiods, longer durations of light were inhibitory towards PSII efficiency in *dm* (Fig 1I). Results here suggest that the smaller *dm* rosette size but not paleness is related to reduced sugar production. Also, the longer durations of light exposure had a detrimental impact on *dm* photosynthesis.

Development-related parameters were further analyzed, since reduced leaf size in *dm* plants pose interesting implications regarding development. Epidermal leaf cell size in *dm* was 57% smaller that WT with cells more densely packed (increased by 52%; Fig 1J,K). A similar evaluation of hypocotyl cell number and length did not show the same effect (Supp Fig 1K,L) indicating that impaired chloroplast-function limits leaf development but not all aboveground tissue. Stomata-based observations reveal that *dm* have a 14% reduced pore length (Fig 1L) but a 30% higher stomatal density (Fig 1M). However, elevated stomatal density was not sufficient to compensate for a smaller stomata pore since *dm* exhibit reduced transpiration rates (Supp Fig 1G). We also evaluated yield-related parameters, which were 10-20% lower overall in *dm* showing reduced silique number, silique length (both by 11-12%), silique pigment levels, inflorescence height (by 17%), and carpel seed number (by 20%; Supp Fig 1M-Q). Alterations in development were observed from bolting-related parameters – where the *dm* bolted at a fewer leaf stage (Supp Fig 1R). *cia2*, *cil,* and *dm* all took longer than WT to bolt: on average taking 2.83, 1.66, and 3 days more respectively (Supp Fig 1S). Thus, CIA2 and CIL mediated chloroplast-function governs plant developmental events including leaf development, yield, and flowering.

### Genotype differences highlight transcriptome and metabolome reprogramming

CIA2 and CIL are essential to multifaceted physiological processes (Fig 1), thus we evaluated underlying reprogramming events that might govern these physiological outcomes across genotypes. Since chloroplast-function relies on transcriptional programs to regulate chloroplast- and photosynthesis-associated nuclear genes (*Cp-* & *PhANG*s), we first identified CIA2 and CIL-mediated transcriptional changes by performing RNA-sequencing on WT, *cia2*, *cil*, and *dm* using excised leaf 5 and 6 on day of bolting (Supp File 2). Differential gene expression (DEG) analysis revealed genotype-associated transcriptome differences and a trend of *dm*>*cia2*>*cil*, based on overall DEG numbers (155 to 45 to 18, respectively compared to WT; Fig 2A), resembling the observed phenotypic characterization trend (Fig 1B). All genotypes had greater numbers of repressed genes, including *dm* with 114 repressed, 41 induced (Fig 2A) together indicating the role of CIA2 and CIL as positive regulators of transcript-expression. An overlap of DEGs across genotypes helped identify 6 and 2 common DEGs that were repressed and induced respectively (Fig 2B,C). Compared to DEGs shared across genotypes, more DEGs were unique to *dm*; 95 and 32 DEGs that were repressed and induced respectively – potentially indicating compensatory effects which are potentiated when both CIA2 and CIL are lacking. Amongst the uniquely repressed *dm* DEGs was chloroplast-related transcription factor (TF) *Golden 2-like 1* (*GLK1*; −2.73 log2FC; Fig 2B). Interestingly we also found *GLK1* as a DEGs of *cia2-2 cil-1* from Gawroński et al., 2021 who utilized young leaves for transcriptomic analysis (Supp Fig 2A). Additionally, there were 18 other shared genes – including *rubisco small chain family protein* (*RBCS2B*), and *thylakoid lumen protein* (AT5G52970; Supp Fig 2A). Using our *dm* vs WT DEG comparisons for gene ontology analysis, we evaluated which biological processes are enriched (Fig 2D; Supp File 3). There was significant overrepresentation of CK metabolism, carbohydrate-related metabolic processes (glucose, fructose, starch), photosynthesis, growth, response to hydrogen peroxide, stomatal closure, chloroplast organization, and negative regulation of leaf senescence amongst others (Fig 2D).

**Figure 2:**
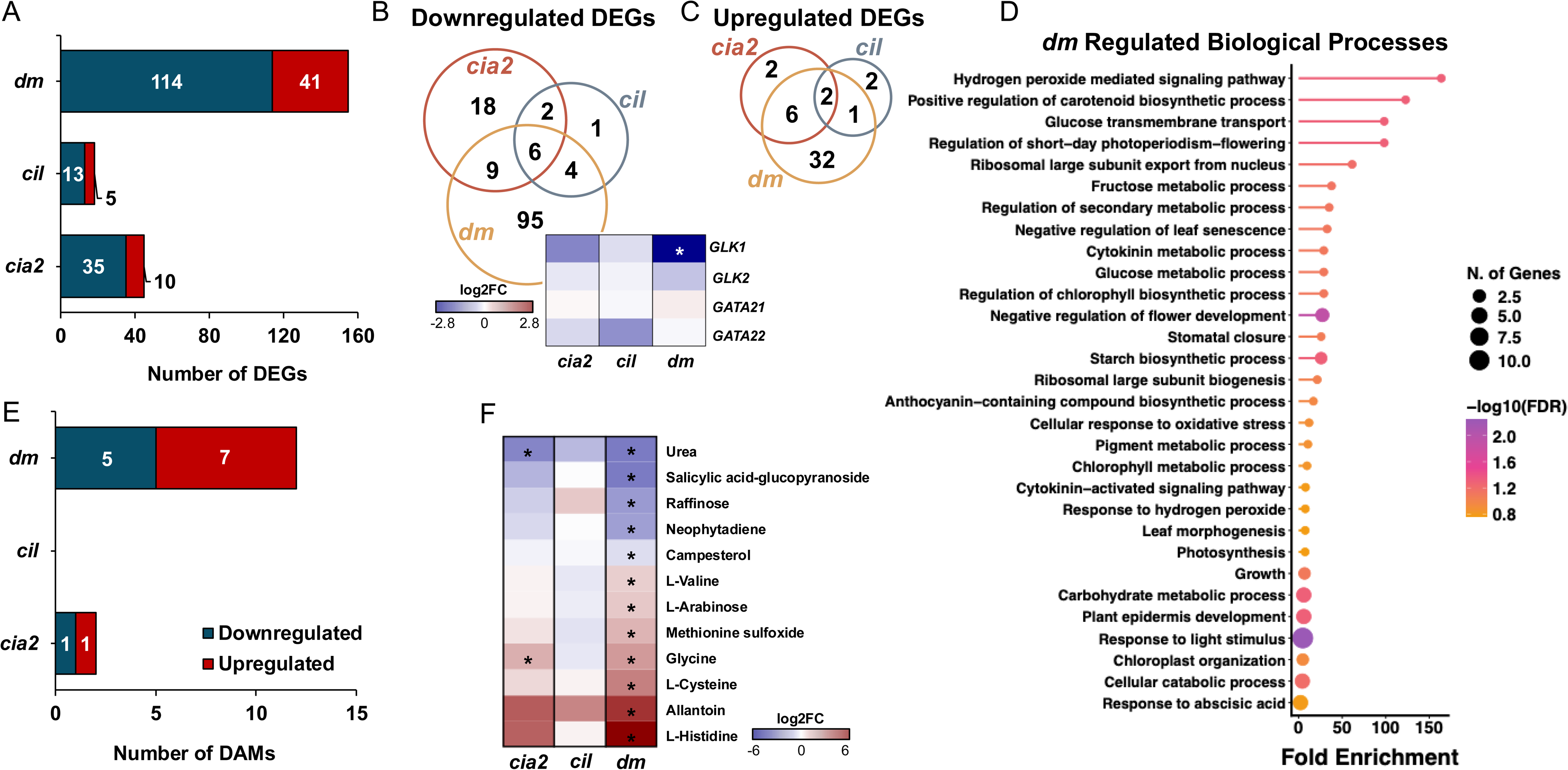
Genotype differences highlight transcriptome and metabolome reprogramming. (A) Number of differentially expressed genes (DEGs) relative to WT (DEG cutoff FDR < 0.05, log2FC = 0) with color used to note significantly upregulated (red) and downregulated (blue) genes. (B-C) Venn diagrams showing unique and shared DEGs that are repressed (B) or induced (C), with numbers noting DEGs. Heatmap represents expression profile of selective genes on a log2FC scale relative to WT with asterisk noting significant difference (FDR < 0.05). (D) Gene ontology biological process analysis of DEGs regulated in *dm* relative to WT. Lollipop plot representing fold enrichment of selective GO terms with dot size used to note number of genes within each GO term category and color noting −log(FDR). (E) Number of differentially abundant metabolites (DAMs) relative to WT (DAM cutoff FDR < 0.05, log2FC = 0) with color used to note significantly upregulated (red) and downregulated (blue) metabolites. (F) Heatmap represents expression profile of DAMs on a log2FC scale relative to WT with asterisk noting significant difference (FDR < 0.05).

Identical to our tissue sampling design for RNA-isolation, we performed metabolite profiling to identify 82 metabolites (>90% confidence) utilized for differentially abundant metabolite (DAM) analysis (Supp File 4). Genotype to WT comparisons revealed a DAM trend resembling the DEG pattern, except for *cil* that showed no DAMs, making the overall DAM trend *dm*>*cia2*>*cil*=WT (Fig 2E). Amongst the 12 DAMs in the *dm*, 5 were repressed while 7 were induced (Fig 2E). Urea showed the strongest repression −3.12 log2FC while L-Histidine was highly induced (6.05 log2FC; Fig 2F). The *dm* also showed a significantly greater abundance of stress-related metabolites including L-Valine (1.15 log2FC; Fig 2F). This suggests that nitrogen and amino acid metabolism are impacted in the *dm*.

### CIA2 and CIL are necessary for cytokinin-mediated chloroplast biogenesis and photosynthesis

Since CK-related biological processes were enriched in the *dm* background (Fig 2D), we evaluated if CIA2 and CIL are needed for CK-mediated chloroplast biogenesis. Dark grown etiolated seedlings were moved to light for 24 h causing de-etiolation or regreening (Fig 3A), during which they were treated ±CK (*trans*-Zeatin; *t*Z; 1 μM), then analyzed for *CIA2* and *CIL* gene expression. After 24 h, both *CIA2* and *CIL* showed induction, with *t*Z treatment further enhancing *CIA2* expression but not *CIL* (Fig 3B). This conforms with previous studies indicating CK induction of *CIA2* and *CIL*: including meta-analysis of CK-responsive genes (Bhargava et al., 2013), eBAR CK-treated (*t*Z 20 μM for 3 h) induction of both *CIA2* and *CIL* (>1.2 log2FC), and leaves treated with *t*Z (1 μM) showing *CIA2* induction (1.13 and 2.46 log2FC by 2 h and 72h) and *CIL* induction (2.49 log2FC by 72 h; Supp Fig 3). Together, this suggests that CK-mediated chloroplast action operates through both *CIA2* and *CIL*, with *CIA2* exhibiting a more robust CK response profile. Further examination of pigment accumulation at 0 h and after 24 h of regreening showed *dm* to have the palest phenotype with reduced chlorophyll a content (Fig 3A, C; Supp Fig 4). *t*Z treatment was able to enhance chlorophyll a content by 44% in WT seedlings, but not in other genotypes (Fig 3C). Chlorophyll b and carotenoid content were not significantly affected by *t*Z (Supp Fig 4). These findings suggest that in *cia2*, *cil*, and *dm*, a CK signal is potentially dampened resulting in a delayed regreening.

**Figure 3:**
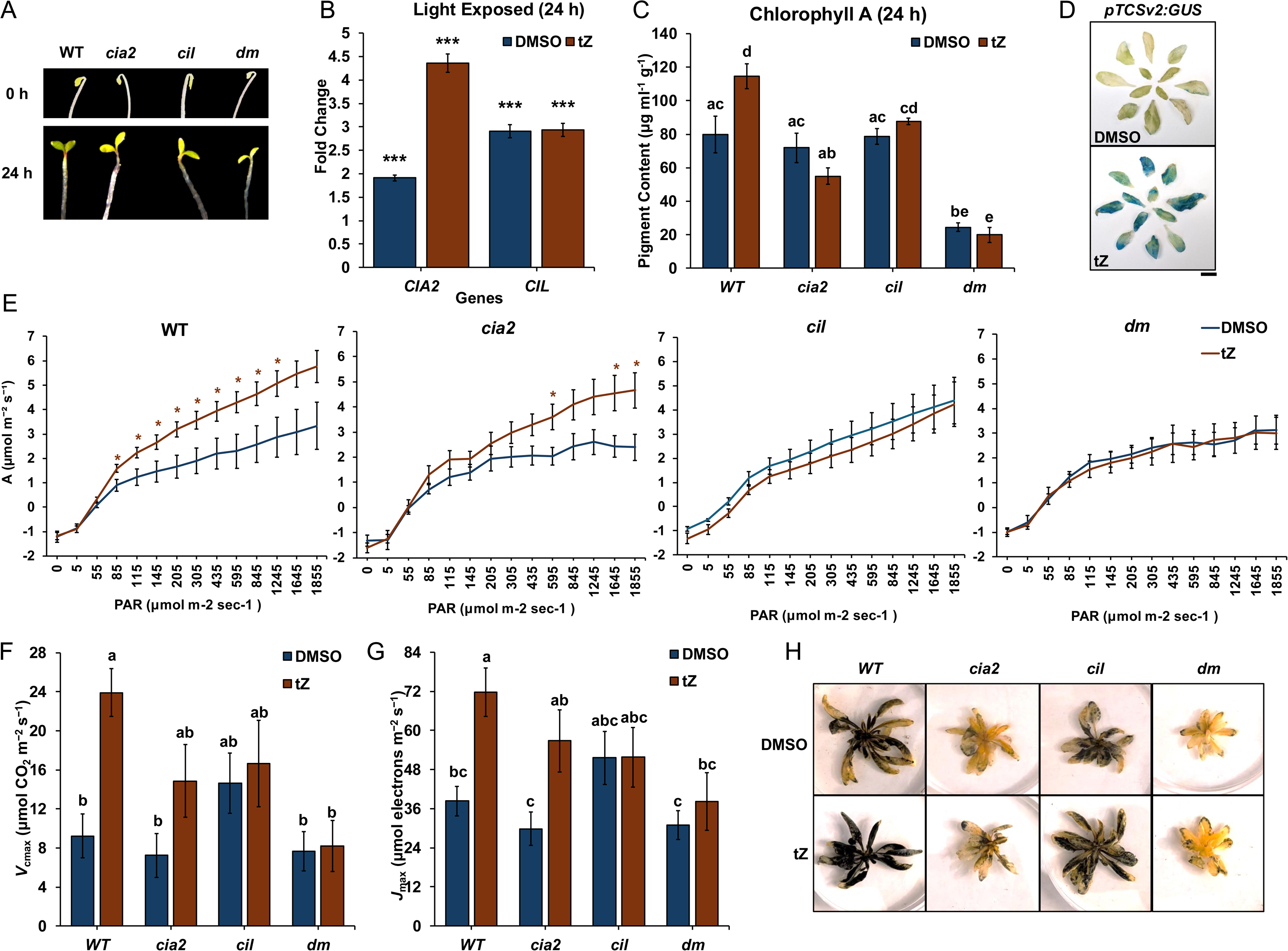
CIA2 and CIL are necessary for cytokinin-mediated chloroplast biogenesis and photosynthesis. (A) Dark-grown seedlings (5 DAG) following or after 24 h of light exposure (representative pictures). (B) qRT-PCR analysis of *CIA2* and *CIL* expression in dark-grown WT seedlings (5 DAG) after 24 h of light exposure under *t*Z (1 μM) or DMSO treatment (noted using colors), with 30 seedlings pooled per sample. Fold change relative to 0 h with *TUB4* as the control primers. (C) Chlorophyll a content in dark-grown seedlings (5 DAG) after 24 h of light exposure under *t*Z (1 μM) or DMSO treatment (noted using colors), with 30 seedlings pooled per sample. (D) *pTCSv2:GUS* (60 DAS) with *t*Z (10 μM) or DMSO treatment were GUS stained and rosette leaves (representative mature/young leaves in outer/inner circle) presented. Scale bar = 1 cm. (E) Genotypes with *t*Z or DMSO treatment were analyzed for photosynthesis (*A*; µmol CO_2_ m^-2^ s^-1^) across photosynthetically active radiation (PAR; µmol m^-2^ s^-1^). Leaf 5-8 were utilized for all LI6800-based photosynthesis analysis using at least 5 plants measured on 60 DAS. (F-G) Maximum rate of rubisco carboxylation (*V_cmax_*; μmol CO_2_ m^−2^ s^−1^) and electron transport (*J_max_*; μmol electrons m^−2^ s^−1^) measured, values normalized to 25 °C before presenting. Statistical test performed using ANOVA and Duncan’s MRT with different letters noting significant difference (p.adj < 0.05). (H) Starch staining with Lugol’s stain at end of night (30 DAS; representative photographs). All results are represented as mean ±SE for at least three biological replicates. Unless noted otherwise, statistical test performed using ANOVA and Tukey’s HSD with different letters noting significant difference (p.adj < 0.05) or asterisk (* p.adj < 0.05, *** p.adj < 0.005).

Since CK prolongs photosynthesis, we watered all genotypes with *t*Z (10 μM) every 5 days starting at 20 DAS, then measured rates of photosynthesis in mature plants (60 DAS). Initially we utilized *pTCSv2:GUS* lines (Steiner et al., 2020) to evaluate if *t*Z watering induces a cytokinin-signaling (two-component signaling; TCS) response and found strong reporter induction across the rosette leaves (Fig 3D), highlighting *t*Z uptake and induction of TCS signal. We hypothesized that TCS components act upstream of CIA2/CIL, since *CIA2*/*CIL* promoters containing several type-B ARR (RRB) binding sites (Supp Fig 5) and due to previous reports utilizing ChIP-sequencing indicating *CIA2* (albeit not *CIL*) as a candidate target of RRBs, ±CK treatment (Zubo et al., 2017, Xie at al., 2018). As such, when TCS signal is induced by *t*Z watering, CIA2/CIL should mediate this signal, leading to an elevated photosynthesis response. We evaluated photosynthesis (*A*) rates across increasing PAR intensities and found that *t*Z significantly elevated photosynthesis levels in WT and *cia2*, however, not in *cil* and *dm* (Fig 3E). *t*Z did not significantly alter *I_50_* and *I_comp_*levels in any genotype, indicating that *t*Z alters photosynthesis beyond the initial 50% saturation (Supp Fig 6A,B). Using *AC_i_* curve analysis, we tested if *t*Z retains higher rates of carboxylation and electron transport. Resembling photosynthesis light curve trends, *t*Z maintained significantly higher *V_cmax_* and *J_max_* levels in WT, by an enormous increase of 159% and 87% respectively relative to DMSO (Fig 3F,G). However, this trend was not observed in the other genotypes, except for *J_max_* in *cia2* (Fig 3F,G). Next, rosettes were stained for starch accumulation at the end of dark and light periods as separate measures of efficient photosynthate remobilization and photosynthesis respectively (Fig 3H and Supp Fig 6C). At end of night, *cia2* and *dm* lacked starch while WT and *cil* still had starch accumulation (Fig 3H). *t*Z was able to elevate starch accumulation at end of night in WT while this effect was less pronounced in *cia2* and *dm* (Fig 3H). Expectedly, at the end of day, all genotypes regardless of treatment showed high starch accumulation (Supp Fig 6C). However, in *dm* the younger leaves had reduced starch accumulation relative to older leaves (Supp Fig 6C). Together, CIA2 and CIL maintain CK activity towards de-etiolation and photosynthesis.

### CIA2 and CIL mediate cytokinin action in senescence

Given that *CIA2* and *CIL* are essential for CK action in chloroplast-function (Fig 3), we predicted they mediate the CK signal to block senescence and performed time course dark-induced senescence (DIS) assays to test this (Fig 4). The initial 0 h timepoint measurement marked >0.83 F_v_/F_m_ across all genotypes as well as a reduction in F_v_/F_m_ as DIS progressed albeit at different rates (Fig 4A-C, Supp Fig 7A). Specifically, *dm* without treatment showed accelerated leaf senescence with a 54% reduction in F_v_/F_m_ relative to WT at 144 h (Fig 4C). This was verified using an independent *cia2-2 cil-1 dm* line revealing a similar reduction (72%) in F_v_/F_m_ relative to WT at 144 h (Supp Fig 8B) – indicating that CIA2 and CIL are necessary for normal progression of senescence.

**Figure 4:**
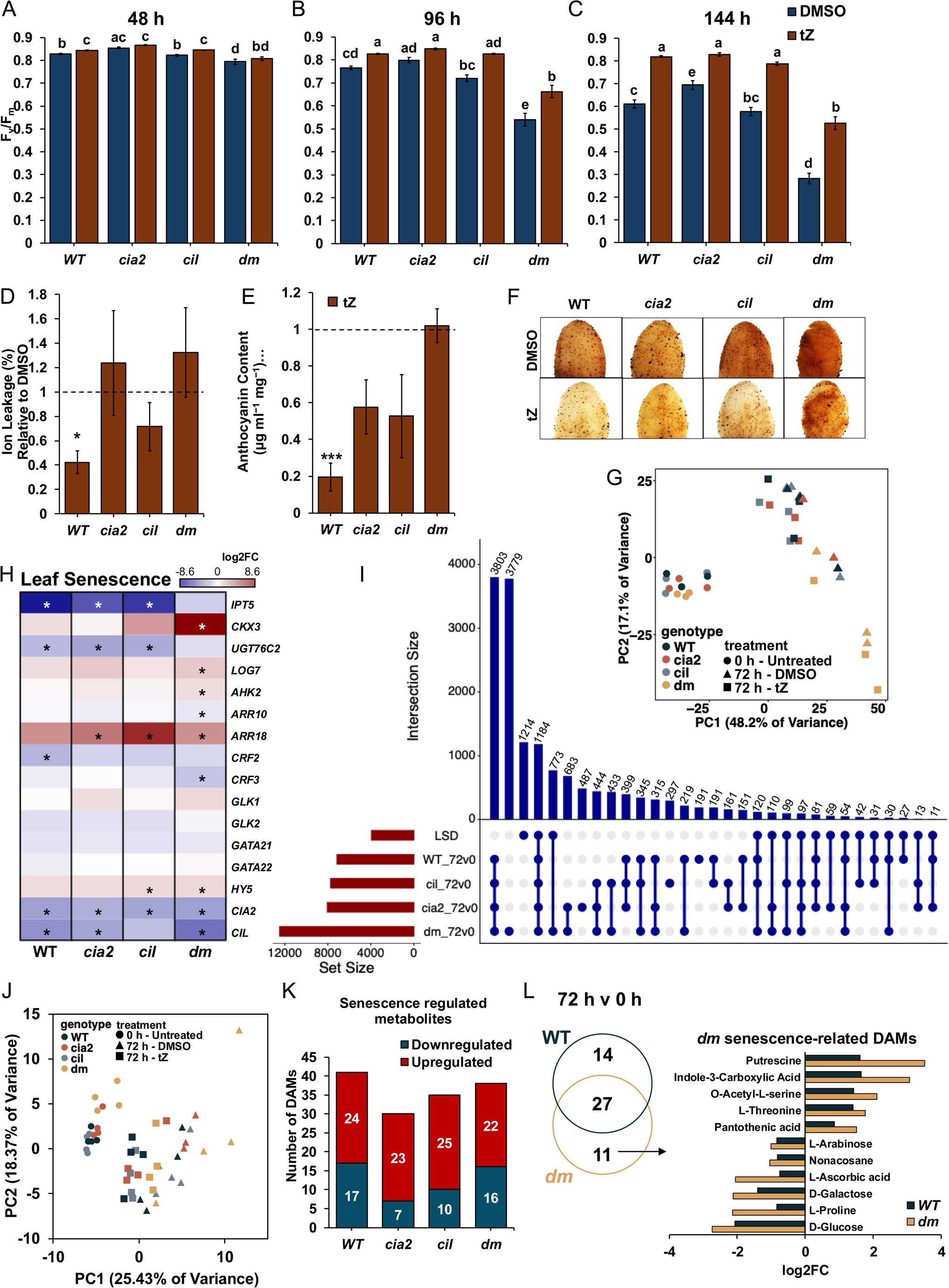
CIA2 and CIL mediate cytokinin action in senescence. (A-C) Photosystem II efficiency (F_v_/F_m_) analysis using leaves 3-8 under dark-induced senescence (DIS) with *t*Z (1 μM) or DMSO treatment (noted using colors). 21-31 leaves were measured per genotype and treatment. (D) Ion leakage, (E) anthocyanin content, (F) and H_2_O_2_ accumulation (3,3’-Diaminobenzidine; DAB staining) analysis of leaves 7-8 following DIS (144 h) under *t*Z (1 μM) or DMSO (dashed line) treatment. For ion leakage and anthocyanin content, at least 9 and 6 leaves respectively were measured per genotype and treatment. All results represented as mean ±SE for at least three biological replicates. Statistical test performed using ANOVA and Tukey’s HSD with different letters noting statistically significant difference (p.adj < 0.05) or asterisk (* p.adj < 0.05, *** p.adj < 0.005). (G) Overall gene expression differences noted using principal component analysis (PCA). PCA was performed for genotypes (leaf 5-6) under untreated (0 h) or DIS 72 h treated with *t*Z (1 μM) or DMSO. Genotypes noted with color while treatment noted with shape for each sample across three biological replicates. (H) Heatmap represents expression profile of selective CK-related genes. Gene expression on a log2FC scale (72 h vs 0 h comparison) with red/blue being induction/repression. Asterisk noting statistically significant difference (FDR < 0.05). (I) Upset plot denoting distinct and shared differentially expressed genes (DEGs) under senescence (72 h vs 0 h). Overall DEGs (FDR < 0.05) along with known senescence-associated genes (SAGs) from leaf senescence database (LSD) were utilized. Brown bars represent total DEGs while blue bars indicate specific intersection (numbers indicate intersection size). (J) Overall metabolite abundance differences noted using PCA, under untreated (0 h) or DIS 72 h treated with *t*Z (1 μM) or DMSO. Genotypes noted with color while treatment noted with shape for each sample across four biological replicates. (K) Differentially abundant metabolites (DAMs; FDR < 0.05) noted with red/blue noting upregulated/downregulated DAMs while numbers indicating total DAMs in each category. (L) Venn diagram noting distinct and shared DAMs across DIS 72 h v 0 h in WT and *dm* genotypes. Bar graph depicts induction and repression trends (log2FC scale) of DAMs distinct to the *dm* under senescence, with color noting WT or *dm*.

Examinations across DIS with CK treatment showed that at 48 h, *t*Z maintained higher F_v_/F_m_ in WT and *cil* but not in *cia2* or *dm* (Fig 4A). However, by 144 h, *t*Z showed retention of F_v_/F_m_ and pigment levels across all genotypes (Fig 4C, Supp Fig 7B, C). Despite this, *t*Z treated *dm* still exhibit significantly reduced F_v_/F_m_ levels compared to *t*Z treated WT (reduced by 35%) indicative of limited CK activity in the *dm* background (Fig 4C). DIS results with independent alleles depict similar trends of reduced *t*Z activity (reduced by 37% in *dm*; Supp Fig 8B). Further, *t*Z significantly elevated chlorophyll a/b ratio in *cia2* and *dm*, while not in WT and *cil* (Supp Fig 7D) indicating that *cia2* and *dm* show a delayed response to *t*Z during DIS. Overall this indicates that CK can maintain PSII efficiency in *dm*, albeit partially, which suggests that prolonged CK exposure (96 – 144 h) can trigger CIA2/CIL independent pathways that act redundantly towards delaying leaf senescence.

We also tested consequences of oxidative stress senescence finding that *dm* showed greater sensitivity to exogenous H_2_O_2_ under DIS (72 h), exhibiting 14% reduced F_v_/F_m_ (relative to 0 h; Supp Fig 7E). Further, ion leakage, anthocyanin content, and H_2_O_2_ distribution under DIS ±*t*Z were evaluated (Fig 4D-F). At 144 h, *t*Z reduced ion leakage 58% and anthocyanin content 81% in WT (respectively), but not significantly in other genotypes (Fig 4E,F). Next, visualization of H_2_O_2_ accumulation revealed that *t*Z reduced levels in WT, *cia2*, and *cil*, however only partially in *dm* (Fig 4F). These results suggest that loss of CIA2 and CIL function limits CK antioxidant action.

Senescence causes transcript and metabolite reprogramming, which was evaluated by RNA-seq analysis and metabolite profiling under DIS at 72 h, since CK effects are most apparent between 48-96 h of DIS (Fig 4A,B). This timepoint also captures omics-based changes leading up to the observed physiological changes at late senescence stages (Fig 4C). Overall transcriptome-based differences were evaluated across 0 h and 72 h using PCA analysis (±*t*Z 1 μM; Fig 4G). Timepoint differences explained majority of the variation during DIS (Fig 4G), indicating a dramatic senescence-based transcriptome reprogramming. A genotype-effect was also evident with *dm* showing distinct clustering from other genotypes for 72 h samples, and only a subtle CK effect was observed (Fig 4G). Since *dm* phenotypically showed accelerated leaf senescence (Fig 4C), we predicted that 72 h vs 0 h comparisons should reveal distinct senescence-based transcriptome patterns. This was seen with overall numbers of DEGs with genotypes following a *dm*>*cia2*>*cil*>WT trend of 12523, 8075, 7760, 7166 respectively (Fig 4I). Senescence-specific transcriptome changes were examined by comparing overlapping senescence-related DEGs to known senescence-associated genes (SAGs; leaf senescence database v5.0; Zhao et al., 2024; Fig 4I). *dm* shared 2467 DEGs (63%) while WT shared only 1538 DEGs (39%) of 3943 known SAGs (Fig 4I). *CIA2* and *CIL* genes were repressed under senescence (Fig 4H). Together with the *dm* exhibiting a greater senescence-associated transcript response, this indicates that CIA2 and CIL act as negative regulators of senescence. Since CK delays senescence, we evaluated CK-related gene expression and found CK metabolism and signaling genes as distinctly expressed in the *dm* (Fig 4H). These included *CKX3* (cytokinin degradation) induced in *dm* (8.60 log2FC) while *IPT5* and *UGT76C2* (involved in CK biosynthesis and N-glucosylation respectively) repressed in all genotypes except *dm* (Fig 4H). Together this suggests there may be endogenously reduced levels of CK base forms (like *t*Z) in *dm*, as the reason behind observed accelerated leaf senescence.

We additionally evaluated metabolite abundance differences by PCA and found the greatest variation due to senescence timepoints – again with 0 and 72 h samples clustering distinctly as well as +*t*Z at 72 h exhibited distinct overall metabolite profiles (Fig 4J). While all genotypes exhibited similar numbers of DAMs based on 72 h vs 0 h comparisons (Fig 4K), there were senescence regulated DAMs distinct to *dm* (relative to WT; Fig 4L). These included sugars such as D-Glucose and D-Galactose, antioxidants such as L-Proline, and L-Ascorbic acid (each reduced more than 2 log2FC; Fig 4L). This signifies lower sugar availability and ROS scavenging potential in *dm* during senescence, thus indicative of *dm* exhibiting higher susceptibility to oxidative stress (Supp Fig 7E).

### CIA2 and CIL co-expressed genes are cytokinin-responsive and associated with physiological performance

Weighted gene co-expression network analysis (WGCNA) was performed to identify genes co-expressed in a CK-specific pattern (Fig 5, Supp File 5). The analysis revealed 45 modules, each named by a unique color and overall module expression noted using eigengene values (Supp Fig 9A,B, Supp Fig 10). We then identified modules exhibiting WT CK-treatment responses, lacking in *dm*. These included MEyellowgreen (137 genes), MEdarkgreen (189 genes), and MEroyalblue (211 genes) with average eigengene values after *t*Z treatment in WT vs *dm* of 0.33 vs −0.02, 0.32 vs −0.03, and 0.31 vs −0.03 respectively (Fig 5A). MEgreenyellow (1376 genes) and MEpurple (883 genes) showed *t*Z induction in WT, *cia2*, and *cil*, however were repressed in *dm* (eigengene values of −0.11 and −0.24 respectively; Fig 5A). Across these CK-responsive modules, GO analysis revealed enrichment of sugar-related processes, cell cycle and cell wall associated terms, amino acid metabolism responses, and plant hormone-related terms (Fig 5C; Supp File 3).

**Figure 5:**
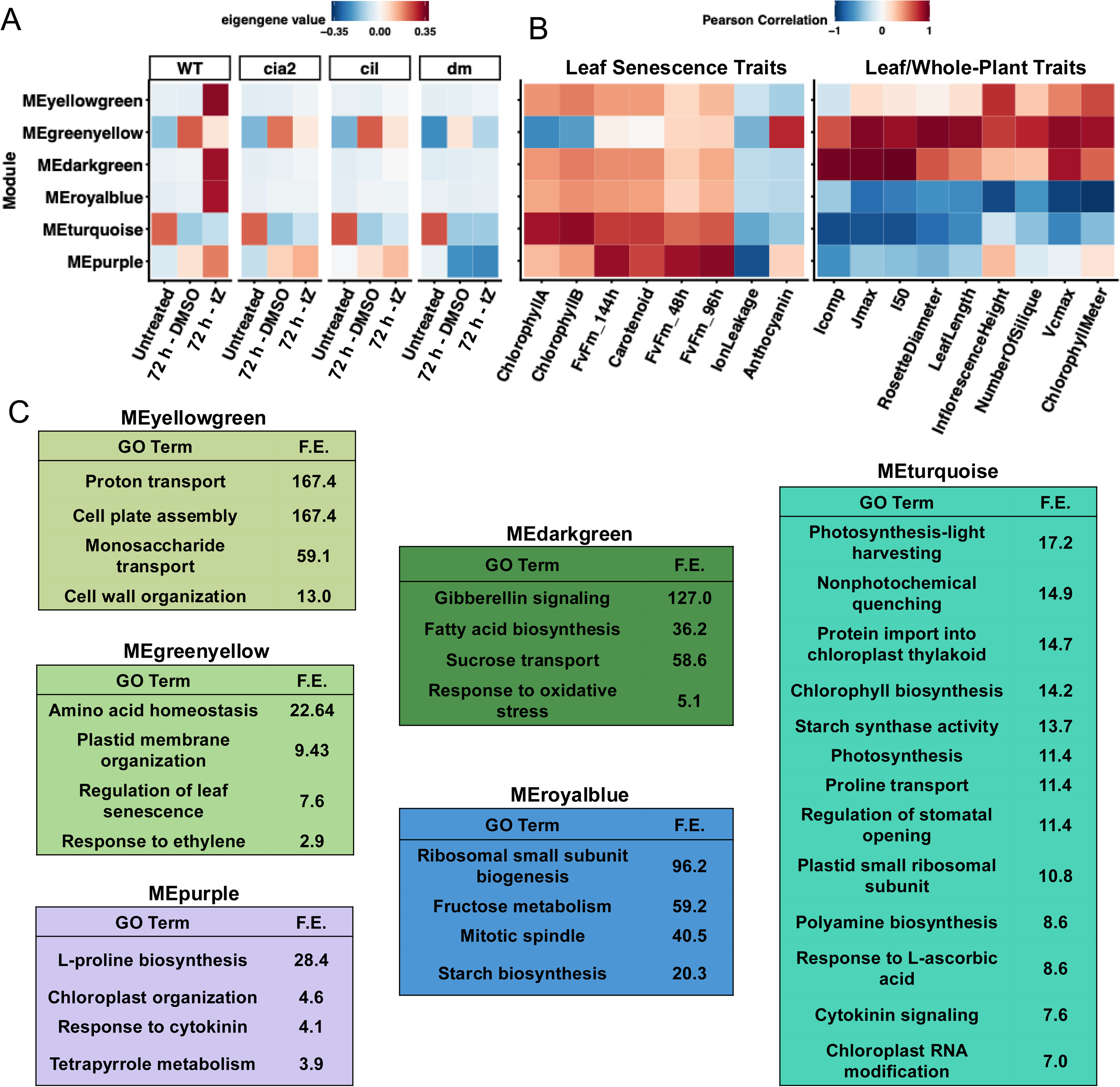
*CIA2* and *CIL* co-expressed genes are cytokinin-responsive and associated with physiological performance. (A) Heatmap representing WGCNA modules (each named with a unique color) eigengene expression (noted with red/blue color gradient) across genotypes in untreated (0 h) and DIS 72 h *t*Z (1 μM) treatment or DMSO conditions. Selective module that show expression changes in a cytokinin-specific manner are depicted. Modules are arranged according to hierarchical clustering. (B) Correlation analysis performed on WGCNA module eigengene expression and physiological traits under DIS (leaf senescence traits) or growth/photosynthesis (leaf or whole-plant traits). Traits clustered on X-axis based on similarity in correlation values. Red and blue colors of the heatmap depict positive or negative correlation (Pearson correlation ranging from 1 to −1). (C) Gene ontology analysis indicating significantly enriched (FDR < 0.05) biological processes across cytokinin-responsive modules. All genes expressed within a module were utilized for GO analysis, following which selective GO terms and their fold enrichment (F.E.) noted.

MEturquoise included the majority of TCS components such as *ARR3, 4, 6, 7, 9, 10, 12, 15, 16, CRF2* as well as CK-metabolism genes like *IPT5* and *CKX4* (Supp File 5). Both *CIA2* and *CIL* were co-expressed in MEturquoise (4454 genes) along with other chloroplast-related transcription factors (TFs) including *GLK2* and *GATA21* (Supp File 5). Correlation analysis of *CIA2* and *CIL* gene expression profiles along with *GLK1,2* and *GATA21,22* revealed *GLK2* and *GATA21,22* to be positively correlated with *CIA2* and *CIL*, while *GLK1* was negative correlated (Supp Fig 11A). The latter is in accordance with *GLK1* being repressed in the *dm* (Fig 2B). Further, *t*Z induced *GATA22* expression occurred in a CIA2-dependent manner, with *cia2* and *dm* lacking this induction (Supp Fig 11B). Overall, this suggests a *CIA2*/*CIL* association with *GLK* and *GATA* factors. Since CIA2 and CIL regulate chloroplast-function, we tested which *Cp-* & *PhANG*s genes were co-expressed, and found this included multiple light-harvesting complex genes (*LHB1B*s*, LHCA*s, *LHCB*s), rubisco subunits (*RBCL, RBCS*s), and ribosomal subunits (*RPL*s*, RPS*s) amongst others (Supp File 5). Amongst the GO terms enriched in MEturquoise were several photosynthesis-associated terms including photosynthesis-light harvesting, nonphotochemical quenching, chlorophyll biosynthetic process, and photosynthesis as well as metabolite-related terms including polyamine biosynthesis process and starch synthase activity (Fig 5C).

Physiological traits under DIS were examined for correlation with WGCNA module expression (eigengene) to reveal associations between CK-responsive gene modules and physiological performance. MEturquoise, MEpurple, MEdarkgreen, MEyellowgreen, and MEroyalblue showed strong positive correlation with PSII efficiency and pigment retention under senescence, with negative correlation to ion leakage (Fig 5B) indicating the association of CK-responsive genes in regulating senescence progression. Since these *t*Z linked modules show enriched terms of chloroplast organization and protein import into the plastid (Fig 5C) their genes may be involved delaying senescence, as is true for the *CIA2* and *CIL* negatively regulating senescence (Fig 4C). Further, correlation of gene module expression with whole-plant growth and photosynthesis traits was performed. MEbrown4, MEplum2, MEivory were amongst the modules showing elevated eigengene expression under untreated (0 h) conditions but not in senescence (72 h) and depicting a strong positive correlation with photosynthesis performance (Supp Fig 10B). Thus, under DIS, genes correlated with photosynthesis performance diminish in expression. WGCNA shows that *CIA2* and *CIL* expression profiles are highly correlated with TCS genes and together are associated with senescence-related physiological performance.

### Cytokinin regulates Cp- & PhANGs through CIA2 and CIL under senescence

Since CIA2 and CIL are known regulators of *Cp-* & *PhANG*s, we evaluated if CK relies on *CIA2* and *CIL* to mediate this action in senescence (Fig 6). Pigment retention from *t*Z treatment was observed during DIS across all genotypes, indicative of a CK-response (Fig 6A). Transcriptional action of *t*Z is evident in DIS with induction of TCS reporter signal and F_v_/F_m_ levels (Supp Fig 12A). Using DEG analysis, we evaluated TCS genes in a *t*Z vs DMSO comparison and found type-A *ARR*s and *AHK4* to be similarly induced across genotypes (Fig 6B). Very minor differences were observed amongst *AHK2*, *ARR2*, *CRF1,* and *CRF5* (Fig 6B, Supp Fig 12B). Similarly, minor transcript differences were observed in CK biosynthesis and degradation genes across genotypes, with a noteworthy exception being *IPT5* that was induced only in WT by *t*Z (Fig 6B, Supp Fig 12C). This suggests that *cia2*, *cil*, and *dm* can show *t*Z TCS induction, ruling out that these genotypes are energetically deficient and thus unresponsive to CK.

**Figure 6:**
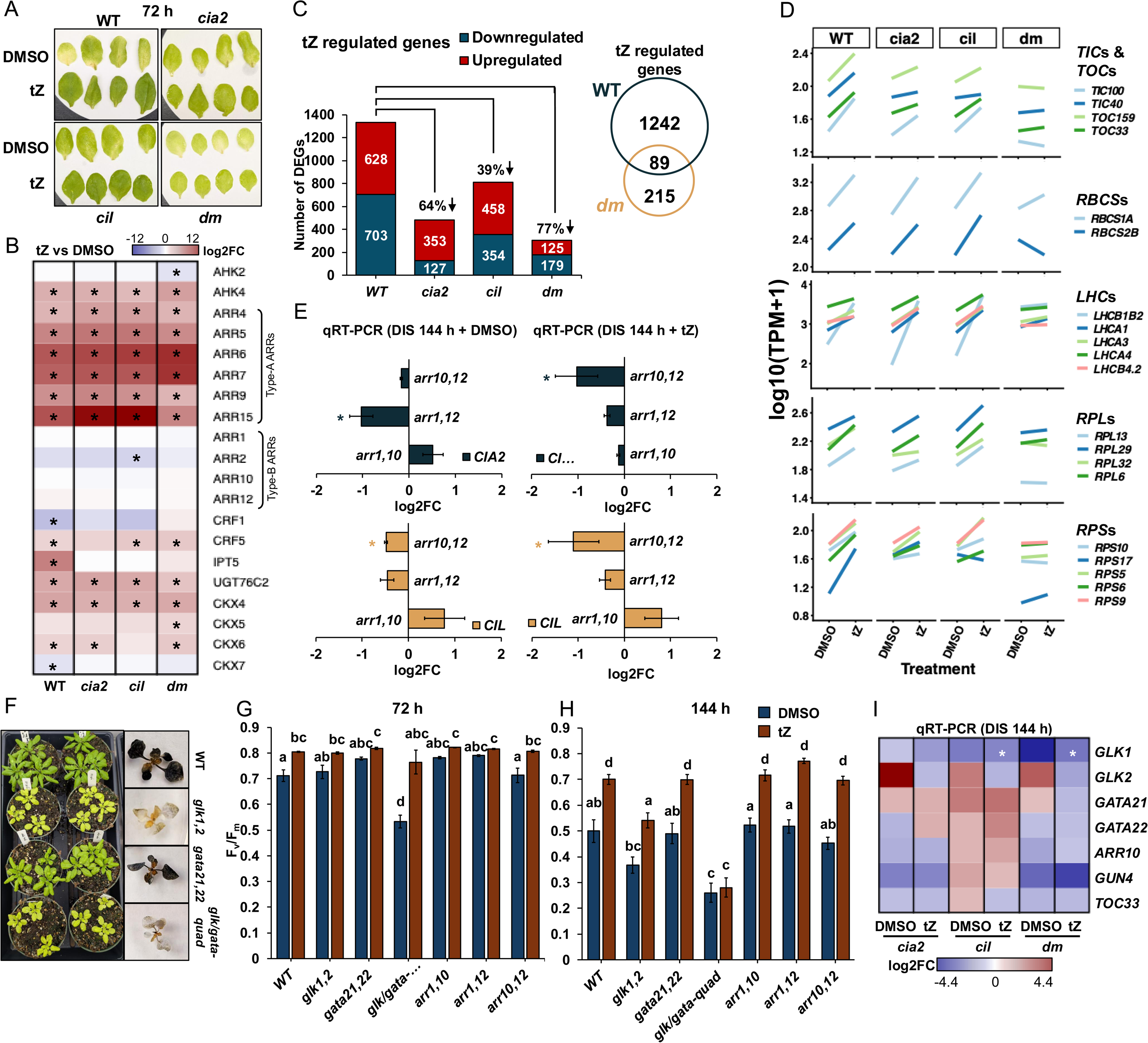
Cytokinin regulates *Cp-* & *PhANG*s through CIA2 and CIL under senescence. (A) Leaves 3-6 following DIS (72 h) under *t*Z or DMSO treatment (representative leaves shown). (B) Heatmap represents expression profile of selective cytokinin-related genes. Gene expression (log2FC scale) in *t*Z vs DMSO with red/blue depicting induction/repression. Asterisk used to note statistically significant difference (FDR < 0.05). (C) DEGs in *t*Z vs DMSO comparison noted with red/blue noting induced/repressed genes while numbers showing DEGs. Numbers above bars indicate percent reduction in total DEGs relative to WT. Venn diagram denoting DEGs in *t*Z vs DMSO for WT or *dm*. (D) Line graphs denoting average gene expression on a log10(TPM + 1) scale under DIS 72 h with *t*Z or DMSO. Color of line denoting a distinct gene as per figure legends. (E) qRT-PCR analysis performed for leaf 5-6 under DIS (144 h) with *t*Z or DMSO. Leaves 5-6 from three plants were pooled for each replicate. Data presented as mean ±SE across three biological replicates, with colors noting *CIA2* or *CIL* gene expression (log2FC). Statistical test performed using Student’s t test with asterisk (* p.val < 0.05) noting significant difference with WT. *TUB4* used as the control gene. (F) Genotypes were grown and photographed on 30 DAS. On 15 DAS, at midday rosettes were detached and stained with Lugol’s stain (representative plants shown). (G-H) PSII efficiency (F_v_/F_m_) analysis measuring leaves 5-6 at DIS timepoints with *t*Z or DMSO (noted using colors). 17-22 leaves were measured (at 72 h 4-6 leaves measured) with results represented as mean ±SE across three biological replicates. Statistical test performed using ANOVA and Tukey’s HSD with different letters noting statistically significant difference (p.adj < 0.05). (I) qRT-PCR analysis performed for leaf 5-6 under DIS (144 h) with *t*Z or DMSO. Leaves 5-6 were pooled from three plants (total 6 leaves) for each replicate and flash frozen. Data presented on log2FC scale with red and blue noting induction and repression respectively across three biological replicates. Statistical test performed using ANOVA and Tukey’s HSD with asterisk (* p.val < 0.05) noting significant difference relative to WT.

We further evaluated if CK signal transduction, downstream of TCS, is affected in *cia2*, *cil*, and *dm*. *t*Z regulated DEGs WT (1331), *cia2* (480), *cil* (812) and *dm* (304; Fig 6C) has a *t*Z responsive trend of WT>*cil*>*cia2*>*dm*, indicating that CIA2 and CIL act together to transduce the CK transcriptional signal. An overlap of *t*Z regulated genes from WT and *dm* revealed 89 commonly regulated genes that show enrichment of CK-related processes (Fig 6C, Supp Fig 12D). Amongst other enriched processes were negative regulation of cell death, aging, and chlorophyll biosynthesis (Supp Fig 12D) – the latter aligning with *t*Z pigment retention results (Fig 6A). PCA of transcriptomic effects in DIS showed that CK-treated samples cluster distinctly, although this effect was least prominent in *dm* (Supp Fig 12E). Overall, this suggests that *dm* is responsive to CK however is limited in CK-signal transduction.

Evaluation of processes that are CK-regulated to delay senescence through CIA2 and CIL was performed by comparing overlap of *t*Z regulated genes, MEturquoise (genes co-expressed with *CIA2* and *CIL*), and genes known to delay senescence (youth-associated genes; YAGs; Guo et al., 2025; Supp Fig 12F). This revealed 74 shared genes whose GO network analysis showed strong association with chloroplast-related complexes (Supp Fig 12F). Thus, *t*Z regulated genes that follow *CIA2* and *CIL* expression patterns are chloroplast-associated and delay senescence. Further, we tested which *Cp-* & *PhANG*s are CK regulated by *CIA2* and *CIL* looking for expression of genes induced by *t*Z in WT but not in *dm*. This revealed several chloroplast-import related genes (*TOC*s/*TIC*s), rubisco-subunit (*RBCS*s), light-harvesting complex genes (*LHC*s), and ribosomal subunits (*RPL*s/*RPS*s; Fig 6D). This highlights that crucial players involved in light-dependent and light-independent reactions, chloroplast import, and chloroplast translation are transcriptionally induced by CK. This effect was evident in WT, *cia2*, and *cil*, however was lacking in *dm*. This indicates that either CIA2 or CIL function can mediate CK transcriptional action, however a loss of CIA2 and CIL function can limit CK-action towards *Cp-* & *PhANG*s under senescence.

Since CIA2 and CIL transduce CK response, we predicted that TCS pathway components regulate *CIA2* and *CIL* expression. Since *RRB*s have binding sites in *CIA2*/*CIL* promoters and potentially target them (Supp Fig 5; Zubo et al., 2017, Xie at al., 2018), we tested if a lack of type-B *ARR* function leads to altered *CIA2* and *CIL* expression under DIS. Utilizing three genotypes: *arr1,10; arr1,12;* and *arr10,12* we found *CIA2* to be repressed in *arr10,12* while *CIL* was repressed in *arr1,12* (Fig 6E). We evaluated if CK-mediated induction of *CIA2* and *CIL* relies on ARR1,10,12 by treating *arr* mutants with *t*Z under DIS, finding *CIA2* and *CIL* to be repressed in *arr10,12* (Fig 6E). This suggests *ARR12* as a potentially regulator of *CIA2* and *CIL*. Since CK-mediated transcriptional regulation of *Cp-* & *PhANG*s operates either directly through *RRB*s or indirectly through *GLK*s/*GATA*s (Cackett et al., 2022) we evaluated *CIA2* and *CIL* expression in: *glk1,2; gata21,22;* and *glk1,2 gata21,22* (hereafter *glk*/*gata*-*quad*). Results showed no significant regulation of *CIA2* and *CIL* under DIS across the *glk*/*gata* genotypes (Supp Fig 12G), implying that *GLK*s and *GATA*s do not rely on the regulation of *CIA2*/*CIL* but regulate chloroplast-function independently.

Testing if *RRB*s or *GLK*/*GATA* show redundancy with *CIA2*/*CIL*, we evaluated type-B *arr* and *glk*/*gata* mutants for growth and senescence-related physiology. All *glk*/*gata* mutants exhibit a pale phenotype, following the trend *glk*/*gata-quad*>*glk1,2*> *gata21,22* (Fig 6F). However, type-B *arr* mutants did not show altered pigmentation (Supp Fig 12H) suggesting redundancy in *RRB* function. *glk1,2* and *glk*/*gata*-*quad* mutants also exhibited reduced starch abundance (Fig 6F). In DIS, the *glk*/*gata*-*quad* resembled *cia2*/*cil*-*dm* by showing accelerated leaf senescence, noted by reduced F_v_/F_m_ at 72 h and 144 h (Fig 6G, H). *t*Z retained F_v_/F_m_ and pigment content in all *arr* and *glk*/*gata* genotypes, albeit limited in the *glk*/*gata*-*quad* mutant, implying that CK can act independent of *GLK1,2* and *GATA21,22* to block senescence (Fig 6G, Supp Fig 12J). Utilizing a previous transcriptome analysis of *glk1,2* (Tu et al., 2022), we found nine shared DEGs to our *cia2*/*cil*-*dm*, including *GLK1* and *RBCS2B*, however genotypes largely show distinct transcriptome regulation (Supp Fig 2B). This was confirmed using qRT-PCR of *cia2*, *cil*, and *cia2*/*cil*-*dm* lines under DIS (±*t*Z) where *GLK1* was significantly repressed in *cil* and *cia2*/*cil*-*dm* by *t*Z indicating that CIA2 and CIL regulates *GLK1* however the reciprocal was not true (Fig 6I, Supp Fig 12G). Overall, *glk*/*gata* resembles *cia2 cil* lines in DIS (Fig 4A-C) indicating that CK can act through *RRB*s on *CIA2*/*CIL* and *GLK*/*GATA* towards prolonging chloroplast-function, however redundancy allows for CK signal to be relayed albeit in a transcriptionally diminished manner.

### Cytokinin regulates leaf senescence through chloroplast metabolism

Abundance of 82 metabolites under untreated (0 h) and DIS (72 h) ±*t*Z, showed distinct patterns of CK-response across genotypes (Fig 7, Supp Fig 13A). Metabolite abundance comparing *t*Z to DMSO revealed 11, 5, and 3 DAMs in *cia2*, *cil*, and *dm* respectively and surprisingly none in WT (Fig 7A, Supp Fig 13B). All DAMs were significant reduced compared to DMSO, however only delta-tocopherol (−4.51 log2FC), L-valine (−1.45 log2FC), and 2-aminobutyric acid (−2.27 log2FC) were significantly repressed in *dm* (Fig 7A). This CK-mediated repression in *dm* indicates a compromised vitamin and amino acid metabolism.

**Figure 7:**
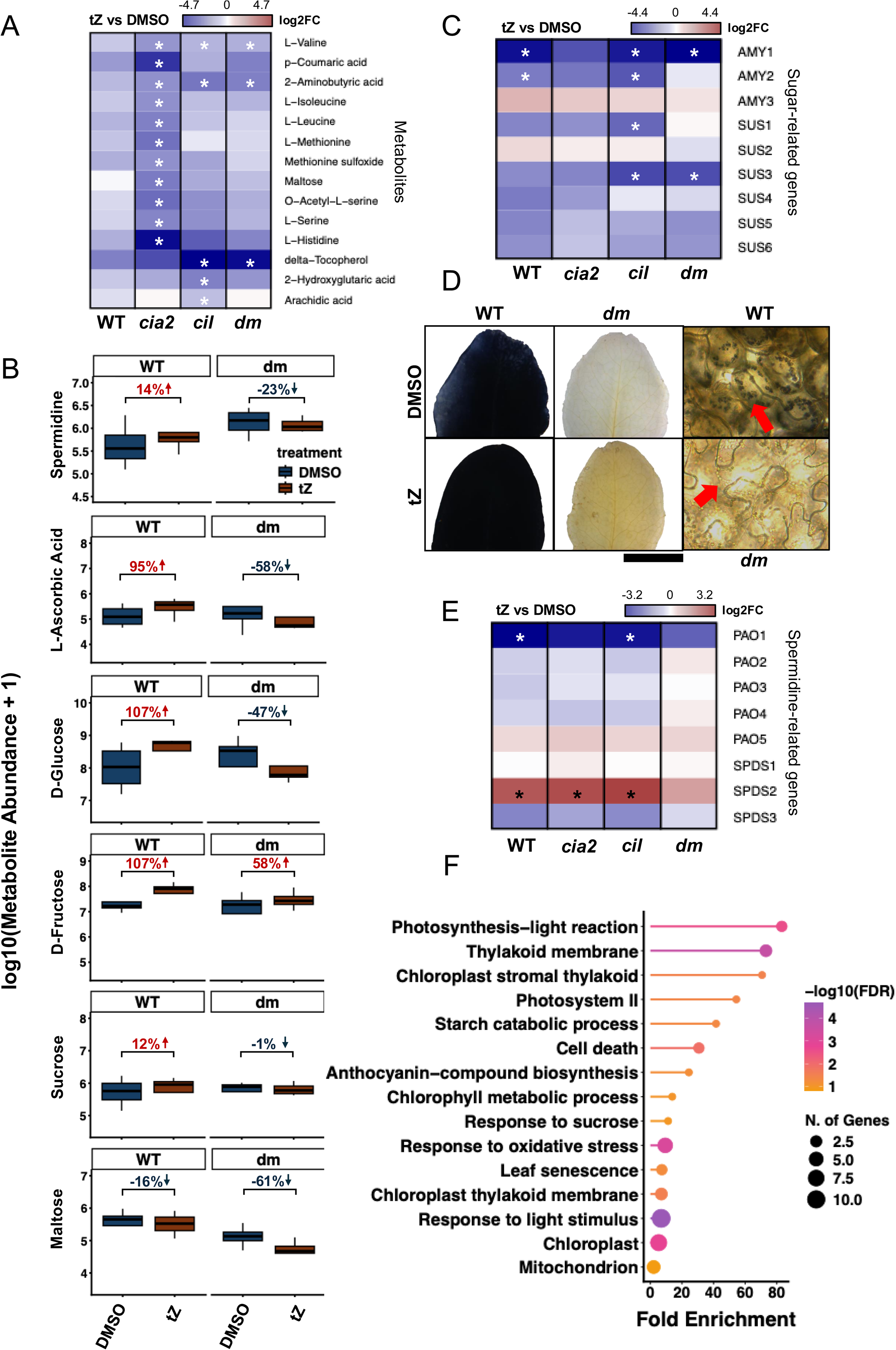
Cytokinin regulates leaf senescence through chloroplast metabolism. (A) Heatmap denotes change in metabolite abundance (log2FC scale) in *t*Z vs DMSO with asterisk noting FDR < 0.05. (B) Box plots used to note average metabolite abundance on a log10(metabolite abundance +1) scale of selective metabolites that show a cytokinin-response in WT but lack similar trends in *dm*. Data presented as average of four biological replicates with numbers indicate change in raw averaged metabolite abundance as a percentage from DMSO to *t*Z treatment (noted with box color), with red/blue text colors indicating increased/decreased percent change. (C,E) Heatmap represents expression profile of sucrose-metabolism (C) and spermidine-metabolism (E) genes. Gene expression represented on a log2FC scale (*t*Z vs DMSO) with red/blue showing induction/repression. Asterisk used to note statistically significant difference (FDR < 0.05). (D) Starch accumulation staining leaf 5-6 from WT or *dm* using Lugol’s stain at DIS 144 h with *t*Z or DMSO (representative photographs shown). Microscope images (40x objective magnification) of *t*Z treated leaves (DIS 144 h), with red arrows pointing to plastids to show stain uptake in WT but not in *dm*. (F) GO analysis of top 100 positively correlated genes with spermidine abundance (determined from integrated transcriptomic-metabolomic analysis). Lollipop plot representing fold enrichment of selective GO terms with dot size used to note number of genes within each GO term category and color noting −log(FDR).

Since *t*Z did not significantly alter the WT metabolite profile, we identified metabolites with distinct CK-responses in WT and *dm* lines based on abundance (Supp Fig 13C). Several metabolites exhibit a CK-mediated repression in *dm* not found to the same extent in WT. These included the sugars (D-glucose, D-fructose, sucrose, and maltose), polyamines such as spermidine, and antioxidants L-ascorbic acid (Fig 7B) – the latter suggesting a greater ROS scavenging response in WT by *t*Z, parallelling H_2_O_2_ distribution findings noted in DIS (Fig 4F). Similarly, in *dm* a reduction of sugar abundance like D-glucose was observed by *t*Z (Fig 7B). Analysis of sugar-related DEGs reveals repression of a sucrose biosynthesis gene *SUS3* by *t*Z in *dm* (−3.03 log2FC) but not in WT (Fig 7C). This trend of *t*Z action was parallelled by examination of DIS leaves with Lugol’s stain, where *t*Z could not retain starch accumulation in *dm* (Fig 7D) although light brown leaf coloration by *t*Z suggested higher α-glucan accumulation (based on Wang et al., 2025). Analysis here highlights a partial activity by CK in *dm* towards resource-maintenance and antioxidants.

Spermidine has been previously shown to regulate the progression of leaf senescence by blocking ethylene-signaling (Liebsch et al., 2022). In our previously reported DIS timeseries analyses, expression of spermidine biosynthesis gene *SPDS2* was significantly induced by *t*Z from 6 to 72h (Khanna et al., 2025; Supp Fig 13D) and from 96 to 144h (Hasannin et al., 2026; Supp Fig 13E). Here, *SPDS2* was also significantly upregulated by *t*Z under DIS in all genotypes except *dm* (Fig 7E). *SPDS2* was also co-expressed with *CIA2* and *CIL* in MEturquoise (Supp File 5). This suggests that loss of CIA2 and CIL function can reduce CK-mediated spermidine biosynthesis under DIS. On the other hand, *SPDS3* (*SPMS*) an enzyme catalyzing spermidine to spermine conversion was repressed across *WT*, *cia2*, and *cil*, although not significantly (Fig 7E). In previous DIS work however, *SPDS3* has been shown to be significantly repressed following *t*Z treatment under DIS (Supp Fig 13D,E). Expression of the genes *PAO2* and *PAO3* catalyzing spermine to spermidine and then to putrescine (Chen et al., 2019), were not significantly regulated (Fig 7E).

We further utilized a mixOmics based modeling approach to integrate our transcriptomic and metabolomic results and identify genes involved in metabolite abundance trends. Amongst genes-metabolites found as highly correlated (correlation cutoff = 0.9) were several chloroplast-related genes including plastid-localized *ALPHA-AMYLASE-LIKE 3* (*AMY3*; involved in starch-metabolism), *FERREDOXIN-NADP(+)-OXIDOREDUCTASE 2* (*FNR2*; a ferredoxin:NADP(H) oxidoreductase present in leaf stroma & thylakoid), *AT4G18810* (*CIA30-type domain protein*), and *FIBRILLIN10* (*FBN10*) along with metabolites including sugars like maltose and antioxidants like L-dehydroacsorbic acid (Supp Fig 14). Further, utilizing integrated transcriptome-metabolome models we evaluated gene profiles showing a strong correlation with spermidine metabolite levels. These top 100 positively correlated genes showed GO enrichment for terms including photosynthesis-light reaction, thylakoid membrane, cell death, response to sucrose, and leaf senescence amongst others (Fig 7F). Overall, this suggests that CK regulation of chloroplast-related metabolites such as spermidine, might be a metabolite-centric mechanism of blocking senescence progression.

## Discussion

Cytokinin (CK) is involved in multifaceted plant responses, including altering chloroplast-function that regulates developmental events such as senescence. CK is known to delay senescence by prolonging photosynthetic performance and pigment content. Specifically, CK treatment activates TCS components that transcriptionally regulate *Cp-* & *PhANG*s. While previous studies described this initial mode of CK action (Kim et al., 2006, Zwack and Rashotte, 2013), little is known about what occurs after primary TCS steps have commenced to bring about dramatic delays in senescence and improved leaf physiology. Here we show chloroplast-import apparatus 2 (CIA2) and CIA2-like (CIL) to be important mediators of CK-action during senescence. These genes were found to be transcriptionally induced by CK in our previous senescence timeseries analysis and here we expand upon their regulation of this process (Khanna et al., 2025). First, we performed phenotypic analyses on loss-of-function *cia2 cil* lines and identified novel physiological functions highlighting CIA2 and CIL to be negative regulators of senescence progression. Then, combining physiological and transcriptomic analyses, we show that CIA2 and CIL are essential for CK-action upon de-etiolation, photosynthesis, and senescence processes. We further show that CK acts through TCS pathway by utilizing *RRB*s towards *CIA2* and *CIL*. Finally, we identify a metabolite-centric mechanism that CK utilizes to block senescence progression in a CIA2/CIL-dependent manner. Finally, we present a model (Fig. 8) depicting cytokinin activity towards chloroplast-function and senescence operating through CIA2 and CIL.

**Figure 8:**
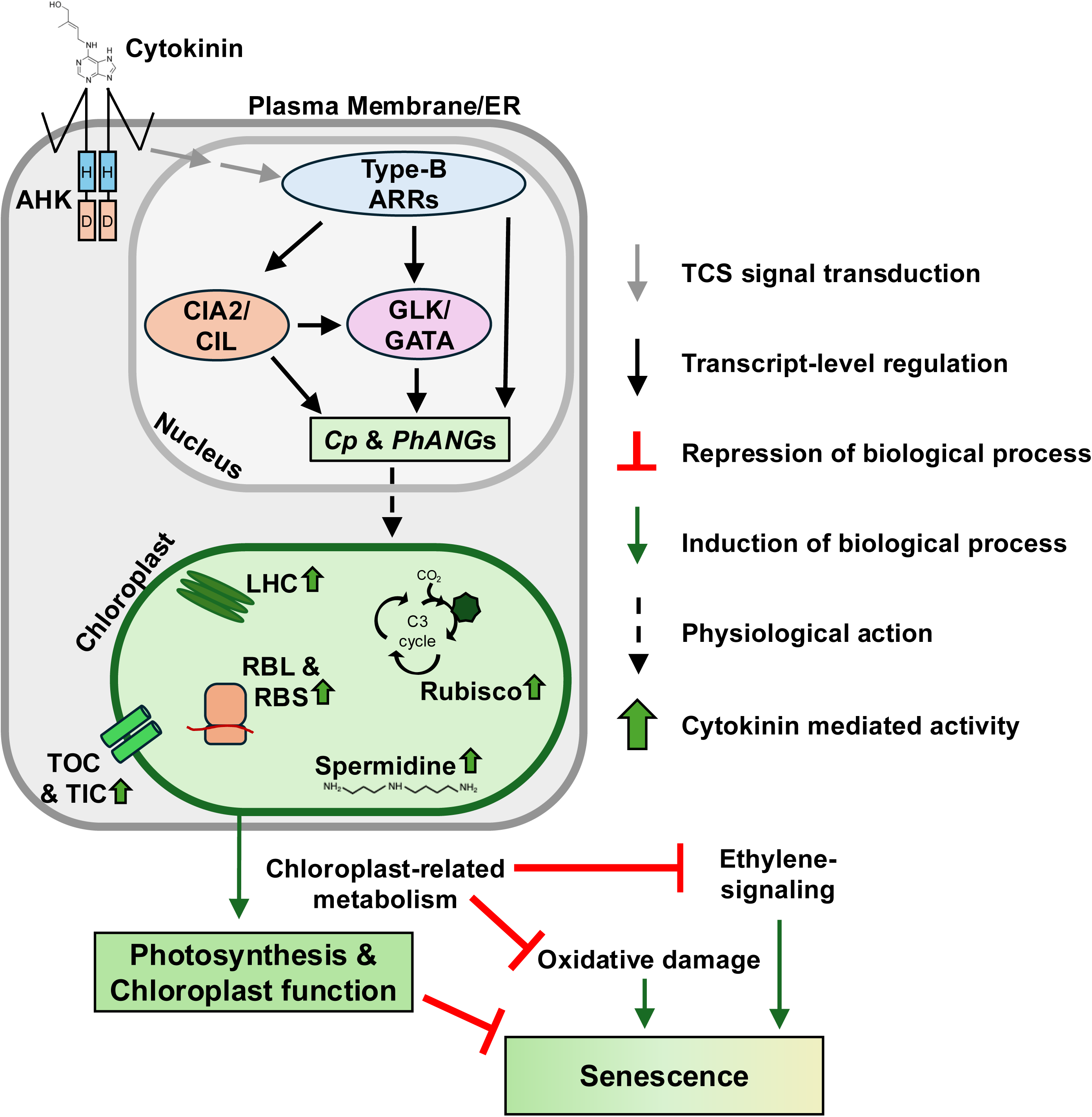
Model of cytokinin activity towards chloroplast-function and senescence operating through CIA2 and CIL. Cytokinin is perceived at the membrane by AHK receptors which initiates a TCS signal that culminates in the activation of type-B ARRs. These activate *Cp-* & *PhANG*s themselves or through GLK/GATA function. Here we propose that cytokinin-mediated activation of *Cp-* & *PhANG*s also operates through CIA2 and CIL. Our work shows that CIA2 and CIL function to regulate *GLK1*, and loss of function in CIA2 and CIL limits cytokinin mediated *TOC*/*TIC*, *LHC*, *RBCS*, *LHC*, and *RBL*/*RBS* along with spermidine metabolite abundance. Through CIA2/CIL mediated transcript and metabolite reprogramming, cytokinin can maintain photosynthesis, limit oxidative damage, and block senescence progression. Specific arrow types noted in figure legends. Model modified from Lim et al., 2007, Cackett et al., 2022, and Liebsch et al., 2022.

### CIA2 and CIL maintain chloroplast-function and rubisco activity

Before we characterized CIA2 and CIL roles as players in CK activity, we performed a detailed phenotypic characterization on *cia2*, *cil*, and their double mutant or *dm* to identify their role in growth and physiology. CIA2 and CIL have previously been shown to alter chloroplast-ultrastructure and function to regulate chloroplast protein-import ability (Li et al., 2021; Gawroński et al., 2021; Yang et al., 2022). Mechanistic insights from previous work indicate these TFs regulate transcriptional output and can bind to *Cp-* & *PhANG*s promoter elements (Sun et al., 2009). Here we expand upon those studies to explore physiological consequences of limiting chloroplast-function by conducting a detailed phenotypic analysis of *cia2*, *cil*, and *dm* lines. We report novel physiological effects of CIA2 and CIL, in additional to consistency in phenotypic trends in prior works i.e. WT=*cil*>*cia2*>*dm* (Li et al., 2021; Gawroński et al., 2021; Yang et al., 2022). Image-based phenotyping identified that loss-of-function in both CIA2 and CIL severely impacts pigment production and growth rates, with CIA2 and CIL acting as redundant partners. These phenotypes are independent of photoperiod, although longer durations of light were detrimental to photosystem efficiency in the *dm*. LiCOR-based light curve analysis revealed that higher light intensities are detrimental to *dm* while NPQ results suggest a reduced utilization of light towards photochemistry in *dm*. CIA2 and CIL were thus essential towards photosynthesis performance. Loss of both CIA2 and CIL function caused depletion of starch reserves, with younger leaves depicting a complete lack of starch accumulation. Application of exogenous sugar was able to recover reduced growth in *dm* to WT levels suggesting limitations in carbohydrate metabolism from impacted chloroplast-function potentially cause this reduced growth. Beyond growth, *dm* constraints leaf development exhibiting reduced leaf size with smaller epidermal leaf cells, and reduced stomata pore lengths. CIA2 and CIL are necessary to maintain normal yield and transition to bolting, the latter being consistent with a separate *cia2-1 cil-3* alleles (Yang et al., 2022). Overall, this phenotypic characterization shows that CIA2 and CIL maintain growth, yield, photosynthesis reactions, and sugar accumulation in a functionally redundant manner, and incorporate extrinsic cues including changes in photoperiod and light intensity.

Secondly, a multi-omics approach was used to identify genotype-based effects towards gene expression and metabolite abundance changes. Paralleling our physiological findings, differentially expressed genes (DEGs) and differentially abundant metabolites (DAMs) followed the *dm*>*cia2*>*cil*≥WT genotype effect trend. Transcriptomic findings suggest that CIA2 and CIL are involved in regulation of other chloroplast-related TFs such as *GLK1*. Since the *dm* (this study) and *glk1,2* (Tu et al., 2022) transcriptional regulation profiles exhibited very little overlap, the transcriptional action of CIA2,CIL and GLK1,2 is largely distinct. *dm* exhibiting repression of *GLK1* shows that CIA2 and CIL act to regulate *Cp-* & *PhANG*s in a manner both independent and through GLK1. *RBCS2B* exhibited shared regulation in CIA2,CIL and GLK1,2 thus highlighting a rubisco-centric mode of action for chloroplast-related transcriptional regulators. As previously reported the *cia2-1 cil-3 dm* exhibits limited rubisco small subunit (RbcS) import into the chloroplast (Yang et al., 2022). This along with reduced transcriptional regulation from this study suggests additive effects in *dm*, causing limited rubsico activity, as observed in *AC_i_* curve analysis, where CIA2 and CIL were required to maintain maximum rate of rubisco activity (*V_cmax_*). Further, DAM profiles indicate compromised nitrogen metabolism in *dm*, specifically of urea – a metabolic byproduct of polyamine biosynthesis pathway. Thus, omics analysis here reveals CIA2 and CIL as modulators of transcripts and metabolite responses involved in chloroplast activity, photosynthesis, and amino acid/nitrogen metabolism.

### Omics analysis reflects the role of CIA2 and CIL as negative regulators of leaf senescence

Chloroplast-function is highly altered across senescence, with it undergoing dismantling prior to other organelles (Schippers et al., 2015). Under DIS, an accelerated leaf senescence phenotype is observed for *dm*. Preceding the strong reduction of F_v_/F_m_ at 144 h, we observe a repression of sugars and antioxidants in the *dm* at 72 h, as well as a global shift in transcriptome (12523 DEGs) and a higher *SAG* response. Together this suggests ineffective oxidative stress mitigation and metabolism in *dm* chloroplast-machinery along with gene expression characteristic of early senescence. Resembling the *cia2*/*cil*-*dm*, the *glk1,2,gata21,22* (*glk*/*gata*-*quad*) genotype also exhibited accelerated senescence. This suggests that lack of functioning chloroplast-related TFs involved in regulating *Cp-* & *PhANG*s can alter chloroplast efficiency and overall trajectory of leaf developmental events like senescence.

The highest proportion of CK regulated proteins are found in the chloroplast (Černý et al., 2013). With the *dm* senescing faster, we evaluated how senescence progression alters CK-metabolism. *dm* showed transcriptional reduction for CK-biosynthesis and elevated CK-degradation, that might lead to lower CK-base form (such as *trans*-Zeatin; *t*Z) availability (Fig 4H). With less CK available to delay senescence, an accelerated leaf senescence phenotype is observed for the *dm*. Exogenous CK can partially rescue this as observed with PSII efficiency findings (Fig 4C). The partial action of CK in *dm* suggests redundancy downstream of TCS pathways to delay senescence. Overall, transcript and metabolite changes in *dm* occurring early in senescence lead to an acceleration of senescence progression. Since, *CIA2* and *CIL* were co-expressed with light-harvesting complexes, rubisco subunits, and ribosomal subunits, the observed repression of *CIA2* and *CIL* under senescence might repression downstream *Cp-* & *PhANG*s and therefore block chloroplast-function.

### Cytokinin enhances de-etiolation and photosynthesis in a CIA2/CIL dependent manner

CK accelerates chloroplast biogenesis through TCS (Cortleven et al., 2016). However, components acting downstream of TCS signal facilitating chloroplast biogenesis and maintenance remain less explored. Transcriptional players that relay this CK activity to the chloroplast may act as crucial regulators of physiological output. CIA2 and CIL are promising candidates as they are known regulators of chloroplast development (Zarrin Ghalami et al., 2026). We thus hypothesized that CIA2 and CIL can maintain chloroplast performance and mediate CK action, and tested responses to exogenous CK (*t*Z) to evaluate this. De-etiolation assay results indicated that losing either CIA2 or CIL function limits CK activity towards seedling regreening. Both *CIA2* and *CIL* are induced under regreening, however CK elevated *CIA2* expression further. This along with previous regulation trends (Supp Fig 3), highlights both *CIA2* and *CIL* as CK-responsive, however *CIA2* with a more robust response.

Further, we show that CK activity extends beyond chloroplast biogenesis, to prolong chloroplast maintenance and photosynthesis, as noted through light and *AC_i_* curve analysis in mature plants (60 DAS). CK retains photosynthesis performance, especially noted under higher PAR (Fig 3E), electron transport, and elevated maximum rate of rubisco carboxylation in a CIA2/CIL dependent manner. Further, CK enabled higher starch accumulation at end of night in WT, while this effect was limited in *dm*. First, this physiological analysis exposes functions of CK that are rarely examined, highlighting the roles of CK in both light-dependent and independent phases of photosynthesis as well as retention of photosynthate reserves. Secondly, this suggests CIA2 and CIL to be essential in relaying these CK-responses, together indicating a crucial role of CIA2 and CIL in both seedlings and mature plants; towards mediating a multi-faceted CK-response including biogenesis and photosynthesis.

### Cytokinin-signaling induces CIA2 and CIL to delay senescence progression

Resembling chloroplast biogenesis and photosynthesis, CIA2 and CIL are involved in CK-response under senescence. *t*Z treatment fails to maintain PSII efficiency levels at early (48 h) senescence timepoints in *cia2* and *dm*. Intriguingly, at later senescence timepoints (144 h) CK can retain PSII efficiency, however only partially compared to WT (Fig 4C). While the genotypes show no differences in CK-perception (Fig 6B), the *dm* shows the fewest DEGs after *t*Z treatment (Fig 6C), indicating a reduced transcriptional output by CK in *dm*. Additionally, the chloroplast can serve as a major source of ROS, but CK application limits H_2_O_2_ accumulation and membrane damage and in a CIA2 and CIL dependent manner. Thus, CIA2 and CIL relay the CK signal induced by exogenous *t*Z treatment and regulate transcriptional output leading to prolonged chloroplast photosynthesis and protection in order to delay senescence progression.

Evaluating the transcriptome findings, we found that CK mediated induction of *Cp-* & *PhANG*s relies on CIA2 and CIL to be functional. CK-action operating through CIA2 and CIL controls expression profiles of *TIC/TOC* transporters, *LHC*s, chloroplast-related ribosomal units (*RBL*s and *RBS*s), and rubisco subunits (*RBCS*) – the latter parallelling maximum rate of rubisco carboxylation findings (Fig 3F). First, this indicates that CK relies on both CIA2 and CIL to mediate a multifaceted chloroplast-functionality response; inducing chloroplast-import, translation, and photosynthesis. Moreover that this response can be relayed if either CIA2 or CIL are functional. Second, these transcriptional profiles were largely unique to CIA2 and CIL, since they regulate a distinct set of transcript targets from what is regulated by GLK1/2 (Supp Fig 2B) but also as they repress *GLK1* (Fig 6I) potentially altering a broad set of *Cp-* & *PhANG*s. Further, *CIA2* and *CIL* are co-expressed with several TCS components including *ARR10,12* along with *GLK2* and *GATA21*. Together these genes are involved in biological processes related to photosynthesis, starch synthesis, antioxidant responses, and spermidine biosynthesis. These also regulated chloroplast-associated genes involved in delaying senescence (youth-associated genes; Guo et al., 2025). Overall, this indicates that CK-action through CIA2 and CIL alters chloroplast metabolism and blocks senescence.

Our testing if CK regulation of *CIA2* and *CIL* operates through RRBs and a GLK/GATA pathway revealed *ARR10,12* as involved in *CIA2* and *CIL* induction by CK, with *ARR12* playing a predominant role (Fig 6E). Although Tu et al., 2022 previously showed GLK1 and GLK2 to target CIA2 and CIL using ChIP-Seq, neither Tu et al., 2022 nor this work (Supp Fig 12G) observed *CIA2* or *CIL* regulation under *glk1,2* or *glk*/*gata*-*quad* mutants based on gene expression analyses. This suggests an *RRB* dependent CIA2 and CIL mode of CK-action, acting on *Cp-* & *PhANG*s, while also on *GLK1*, to control chloroplast-function and delay senescence (Fig 8). Together, loss of CIA2 and CIL does not impact CK perception, however it limits CK signal transduction. This cascade, regulated by *RRB*s, in turn modulates *Cp-* & *PhANG*s transcription causing a compromised response to exogenous CK in *dm*.

### Cytokinin mediated spermidine accumulation represses senescence

Beyond prolonging chloroplast-associated transcriptional regulation, metabolite profiles indicate how CK negatively regulates senescence. CK has previously been shown to alter polyamine levels in *Arabidopsis* (Černý et al., 2013). Spermidine is a polyamine abundant in the plastid that can bind to chloroplast thylakoid membranes under DIS (in barley) and block ethylene signaling components (*EIN3*) to delay leaf senescence (Sobieszczuk-Nowicka et al., 2015, Fürtauer et al., 2019, Liebsch et al., 2022). DIS time series assays have previously shown robust induction of spermidine biosynthesis genes by *t*Z (Supp Fig 13D,E). Parallelling these findings, here *t*Z application induced spermidine biosynthesis genes in all genotypes except *dm*. Beyond transcriptional regulation, CK application can elevate spermidine metabolite content in DIS, however this accumulation occurs late into senescence (7-10 days in barley leaves; Sobieszczuk-Nowicka et al., 2016). Our examination of spermidine levels at a relatively earlier DIS timepoint (72 h) found that application of *t*Z elevates spermidine abundance in WT however reduced its levels in *dm*. While the *dm* depicted already high spermidine levels, this trend is suggestive of higher spermidine accumulation at later DIS timepoints in WT, leading to retention of physiological efficiency. However, since this work did not observe late DIS metabolite changes, whether spermidine accumulation fails to prolong senescence in *dm* remains untested. To evaluate spermidine action in senescence, we utilized an integrated omics approach and tested which genes are associated with spermidine levels. This analysis revealed leaf senescence, autophagy, and photosynthesis related genes as connected. Thus, CK elevates spermidine accumulation in a CIA2/CIL dependent manner that prolongs photosynthesis and regulates autophagy – suggesting a novel mechanism through which CK can repressing senescence progression.

## Methods

### Plant Growth Conditions

*Arabidopsis thaliana* (Arabidopsis) seeds were procured from ABRC (Ohio State University, USA) all being in Col-0 background. Seeds were sown in pots of Promix BX planting mix and grown in 16 h/8 h light (100 μmol m^−2^ s^−1^)/dark cycle with 22 °C/18 °C light/dark temperatures. *cia2-1* (CS6522) was previously described in Sun et al., 2001. Li et al., 2021 developed a crispr based knock-out allele of *cil* (CS2110093) and crossed with *cia2-1*, generating a double mutant (CS2110094). Since these remain unnamed, we proceeded to call them *cil-5*, and *cia2-1 cil-5* in this study – hereafter *cia2-1*, *cil-5*, and *cia2-1 cil-5* being referred as *cia2*, *cil*, and *dm* respectively. A second set of alleles previously described in Gawroński et al., 2020 - *cia2-2* (SALK_004037) and *cil-1* (SAIL_228_C01) were obtained and crossed to develop an independent *cia2-2 cil-1* double mutant. Other genotypes utilized in the study included: *glk1,2* (CS72322), *gata21,22* (CS71765), *glk1,2 gata21,22* (CS71766; hereafter *glk*/*gata*-*quad*), *arr1,10* (CS39990), *arr1,12* (CS6981), and *arr10,12* (CS39991). The *pTCSv2:GUS* seeds were a generous gift from Dr. Maya Bar, Ben Gurion University, Israel.

An overview of sampling approaches utilized in this study is described in Supp Fig 15. Refer to Supp File Methods for a detailed account of materials and methods utilized in this work.

### Image based phenotyping, growth- and development-related parameters

Raspberry Pi cameras-based imaging along with ImageJ analysis was performed for rosette and leaf growth rates. For bolting analysis, seeds were kept in 4 °C for 72 h to stratify before sowing on planting mix following which yield-related parameters recorded on 80 DAS. For cell size and stomatal parameters, epidermal peels were prepared from leaf 5 and visualized using Nikon Eclipse 80i microscope. Refer to Supp File Methods for details.

### Leaf senescence assay and physiology-related parameters

On the day of bolting, leaves were excised and utilized for dark-induced leaf senescence (DIS) assay as per Johnston et al., 2026. For photosystem II efficiency analysis (F_v_/F_m_), measurements were recorded using Open FluorCam (Photon Systems Instruments). For pigment analysis (leaf 3,4), chlorophyll A, chlorophyll B, and carotenoids content was calculated using Sumanta et al., 2014. Anthocyanin concentration (leaf 7,8) was calculated using Lindoo and Caldwell, 1978. Ion leakage analysis (leaf 7,8), was performed resembling Zhang et al., 2020. For H_2_O_2_ histochemical staining (leaf 7,8) the Daudi and O’Brien et al., 2012 protocol was utilized and leaves were imaged using a Nikon SMZ1500 stereomicroscope, and photographed with a Qimaging Fast 1394 digital camera. For RT-PCR analysis, leaf 5,6 were flash frozen after 144 h of DIS and stored in −80 °C until grinding was performed. Refer to Supp File Methods for details.

### Photosynthesis analysis

Photosynthesis and CO_2_ exchange parameters were measured using LI-6800 Portable Photosynthesis System (LICOR biosciences. Lincoln, Nebraska, USA) at 45 DAS or 60 DAS as mentioned in text. Referring to Eckstein et al., 2012 and Pankasem et al., 2024 for light curve and *A*C*_i_* curves analysis respectively the PAR intensities and reference CO_2_ concentrations were used – as noted in Supp File Methods. Analysis of net photosynthetic light-response curve was performed as per Lobo et al., 2013, generating *I_50_* and *I_comp_* using the model described in Kaipiainen, 2009. Using *AC_i_* curves, we calculated maximum rate of rubisco carboxylation (*V_cmax_*) and rate of electron transport (*J_max_*) as per the model described in Bernacchi et al., 2001, 2003. Measurements were corrected to 25 °C before plotting. Refer to Supp File Methods for details.

### Physiological analysis

For starch analysis, soil-grown plants that were *t*Z (10 μM) or DMSO watered were sampled at 30 DAS for starch distribution analysis using Lugol’s stain. For sucrose gradient, seeds were stratified and plated on MS 1x with 1% sucrose and later moved to 0%, 1% and 2% sucrose. After 7 days, ImageJ or F_v_/F_m_ analysis performed. For photoperiod analysis, 5 DAS seeds were moved to short day (8 h/16 h day/night), long day (16 h/8 h day/night), or continuous light conditions (24 h day) for 7 additional days and F_v_/F_m_ recorded. For de-etiolation assay, seeds were kept in dark for 7 days following which they were transferred for 24 h to 3 mM MES containing *t*Z (1 μM) or DMSO in light conditions and utilized for qRT-PCR or pigment analysis. Refer to Supp File Methods for details.

### Transcriptomic analysis

Leaf 5-6 were excised on day of bolting, and flash frozen in liquid nitrogen and stored at −80°C until grinding. Total RNA was isolated via RNeasy kit (Invitrogen) followed by illumina 150×2 sequencing at Novogene Corporation Inc. The following bioinformatic programs were utilized as part of genome assemble and transcript analyses: FastQC (Andrews, 2010), Trimmomatic (Bolger et al., 2014), HiSAT2 and StringTie (Pertea et al., 2016), DeSEQ2 (Love et al., 2014), gene ontology (GO) analysis (Ge et al., 2020), with TAIR 10 *Arabidopsis thaliana* genome (Araport11 annotation) used as the reference genome. Resembling the analysis performed in Khanna et al., 2025, WGCNA package was utilized to identify co-expressed gene modules (Langfelder and Horvath, 2008). Raw transcriptome data is available under Bioproject ID: PRJNA1469230. Refer to Supp File Methods for details.

### Metabolomic analysis

Paralleling the transcriptomic analysis, leaf 5-6 across all four genotypes were excised and lyophilized until metabolite extraction was performed across four biological replicates. Metabolomics samples were extracted using a methyl tert-butyl ether/methanol mixture across four biological replicates, following protocols described previously (Dufková et al., 2023, Berková et al., 2026) with minor modifications. Differentially abundant metabolites reported after pairwise differential abundance analyses performed using the limma R package (version 3.66.0; Ritchie et al., 2015). Refer to Supp File Methods for details.

### Integrated transcriptomic-metabolomics analysis

RNA-sequencing and metabolomic analysis were integrated using the mixOmics package (Rohart et al., 2017) utilizing both multiblock sparse partial least squares discriminant analysis (sPLS-DA) or DIABLO model and a global PLS modeling approach following default parameters. Refer to Supp File Methods for details.

## Acknowledgement

The *pTCSv2:GUS* seeds were a generous gift from Dr. Maya Bar, Ben Gurion University, Israel. Bioinformatic analysis scripts utilized in RNA-sequencing analysis were originally obtained from Dr. Tonia Schwartz, Auburn University as part of the course BIOL6800: Functional Genomics and later modified as needed.

## Author contributions

R.R.K. and A.M.R. conceived the idea; R.R.K., S.S., M.B., S.C., and P.H. performed experiments and data collection; R.R.K., A.S., M.Č., and A.M.R. designed experiments and interpreted research findings; R.R.K., and A.M.R. wrote the original draft; A.M.R. supervised the project; all authors contributed to reviewing and editing the manuscript.

## Supplementary Material Information

The following supplementary material is available with this article:

1. Supplementary Figures
2. Supplementary Methods
3. Supplementary File 1: Growth, photosynthesis, and physiological analysis
4. Supplementary File 2: Transcriptomic and DEG analysis
5. Supplementary File 3: Gene ontology analysis
6. Supplementary File 4: Metabolomic and DAM analysis
7. Supplementary File 5: Gene co-expression (WGCNA)
8. Supplementary File 6: List of qRT-PCR primers

## Data Availability

All data are incorporated into this article and its supplementary material. Transcriptome data available under Bioproject ID: PRJNA1469230.

## Declaration of generative AI

During the data analysis portion of this work, the authors used Google Gemini and OpenAI ChatGPT for evaluating bioinformatics scripts and assistance with R based figure preparation or statistical testing. Following the use of generative AI tools, the authors reviewed and modified the scripts or figures as needed and take complete responsibility for the published content that is part of this article.

## References

Andrews, S. (2010). FastQC: A Quality Control Tool for High Throughput Sequence Data. Retrieved from http://www.bioinformatics.babraham.ac.uk/projects/fastqc/

Argueso CT, Kieber JJ. Cytokinin: From autoclaved DNA to two-component signaling. The Plant Cell. 2024 May;36(5):1429–50.

Berková V, Berka M, Kuzmenko M, Koukalová Š, Bílek O, Novák J, Kováč J, Ďurkovič J, Čičmanec P, Kopriva S, Brzobohatý B, Černý M. Altered cytokinin signaling drives plant drought resilience through lipid remodeling, nutrient homeostasis, cell wall reinforcement, and modulation of biotic stress–associated pathways. Plant Physiology and Biochemistry. 2026 May 20:111402.

Bernacchi CJ, Pimentel C, Long SP. In vivo temperature response functions of parameters required to model RuBP-limited photosynthesis. Plant, Cell & Environment. 2003 Sep;26(9):1419–30.

Bernacchi CJ, Singsaas EL, Pimentel CA, Portis Jr AR, Long SP. Improved temperature response functions for models of Rubisco-limited photosynthesis. Plant, cell & environment. 2001 Feb;24(2):253–9.

Bhargava A, Clabaugh I, To JP, Maxwell BB, Chiang YH, Schaller GE, Loraine A, Kieber JJ. Identification of cytokinin-responsive genes using microarray meta-analysis and RNA-Seq in Arabidopsis. Plant Physiology. 2013 May 1;162(1):272–94.

Bolger AM, Lohse M, Usadel B. Trimmomatic: a flexible trimmer for Illumina sequence data. Bioinformatics. 2014 Aug 1;30(15):2114–20.

Breeze E, Harrison E, McHattie S, Hughes L, Hickman R, Hill C, Kiddle S, Kim YS, Penfold CA, Jenkins D, Zhang C. High-resolution temporal profiling of transcripts during Arabidopsis leaf senescence reveals a distinct chronology of processes and regulation. The Plant Cell. 2011 Mar 1;23(3):873–94.

Bresson J, Bieker S, Riester L, Doll J, Zentgraf U. A guideline for leaf senescence analyses: from quantification to physiological and molecular investigations. Journal of Experimental Botany. 2018 Feb 6;69(4):769–86.

Cackett L, Luginbuehl LH, Schreier TB, Lopez-Juez E, Hibberd JM. Chloroplast development in green plant tissues: the interplay between light, hormone, and transcriptional regulation. New Phytologist. 2022 Mar;233(5):2000–16.

Carillo P, Ferrante A. Decoding the intricate metabolic and biochemical changes in plant senescence: a focus on chloroplasts and mitochondria. Annals of Botany. 2025 Jan 30:mcaf003.

Černý M, Kuklová A, Hoehenwarter W, Fragner L, Novák O, Rotková G, Jedelský PL, Žáková K, Šmehilová M, Strnad M, Weckwerth W. Proteome and metabolome profiling of cytokinin action in Arabidopsis identifying both distinct and similar responses to cytokinin down-and up-regulation. Journal of Experimental Botany. 2013 Nov 1;64(14):4193–206.

Chen D, Shao Q, Yin L, Younis A, Zheng B. Polyamine function in plants: metabolism, regulation on development, and roles in abiotic stress responses. Frontiers in plant science. 2019 Jan 10;9:1945.

Chiang YH, Zubo YO, Tapken W, Kim HJ, Lavanway AM, Howard L, Pilon M, Kieber JJ, Schaller GE. Functional characterization of the GATA transcription factors GNC and CGA1 reveals their key role in chloroplast development, growth, and division in Arabidopsis. Plant physiology. 2012 Sep 1;160(1):332–48.

Cortleven A, Marg I, Yamburenko MV, Schlicke H, Hill K, Grimm B, Schaller GE, Schmülling T. Cytokinin regulates the etioplast-chloroplast transition through the two-component signaling system and activation of chloroplast-related genes. Plant Physiology. 2016 Sep 1;172(1):464–78.

Cortleven A, Schmülling T. Regulation of chloroplast development and function by cytokinin. Journal of experimental botany. 2015 Aug 1;66(16):4999–5013.

Dufková H, Berka M, Psota V, Brzobohatý B, Černý M. Environmental impacts on barley grain composition and longevity. Journal of Experimental Botany. 2023 Mar 13;74(5):1609–28.

Eckstein A, Zięba P, Gabryś H. Sugar and light effects on the condition of the photosynthetic apparatus of Arabidopsis thaliana cultured in vitro. Journal of Plant Growth Regulation. 2012 Mar;31(1):90–101.

Fürtauer L, Küstner L, Weckwerth W, Heyer AG, Nägele T. Resolving subcellular plant metabolism. The Plant Journal. 2019 Nov;100(3):438–55.

Gawroński P, Burdiak P, Scharff LB, Mielecki J, Górecka M, Zaborowska M, Leister D, Waszczak C, Karpiński S. CIA2 and CIA2-LIKE are required for optimal photosynthesis and stress responses in Arabidopsis thaliana. The Plant Journal. 2021 Feb;105(3):619–38.

Ge SX, Jung D, Yao R. ShinyGO: a graphical gene-set enrichment tool for animals and plants. Bioinformatics. 2020 Apr 15;36(8):2628–9.

Guo X, Wang Y, Zhao C, Tan C, Yan W, Xiang S, Zhang D, Zhang H, Zhang M, Yang L, Yan M. An Arabidopsis single-nucleus atlas decodes leaf senescence and nutrient allocation. Cell. 2025 May 29;188(11):2856–71.

Guo Y, Ren G, Zhang K, Li Z, Miao Y, Guo H. Leaf senescence: progression, regulation, and application. Molecular horticulture. 2021 Jun 16;1(1):5.

Hallmark HT, Rashotte AM. Review–cytokinin response factors: responding to more than cytokinin. Plant Science. 2019 Dec 1;289:110251.

Hasannin O, Khanna RR, Singh S, Petřík I, Strnad M, Novák O, Černý M, Rashotte AM. Cytokinin senescence delay is shaped by receptor specificity and metabolic stability. Plant Physiology. 2026 Jul;201(3):kiag392.

Johnston GA, Hasannin O, Khanna RR, Rashotte AM, Argueso CT. Root growth and leaf senescence assays as physiological readouts of cytokinin response in Arabidopsis Thaliana. InPlant Hormones: Methods and Protocols 2026 Apr 1 (pp. 247–260). New York, NY: Springer US.

Kábrtová V, Kučerová Z, Budíková B, Hudeček M, Špundová M, Plíhal O. Cytokinin availability and chloroplast functional lifetime in senescing leaves. Frontiers in Plant Science. 2026 Jul 27;17:1901011.

Kaipiainen EL. Parameters of photosynthesis light curve in Salix dasyclados and their changes during the growth season. Russian journal of plant physiology. 2009 Jul;56(4):445–53.

Khanna RR, Hasannin O, Thennakoon M, Buchheit A, Petřík I, Strnad M, Novák O, Černý M, Rashotte AM. trans-Zeatin N-glucosides can delay salt accelerated leaf senescence in Arabidopsis thaliana. Plant physiology and biochemistry. 2025 Oct 1;227:110101.

Kim HJ, Ryu H, Hong SH, Woo HR, Lim PO, Lee IC, Sheen J, Nam HG, Hwang I. Cytokinin-mediated control of leaf longevity by AHK3 through phosphorylation of ARR2 in Arabidopsis. Proceedings of the National Academy of Sciences. 2006 Jan 17;103(3):814–9.

Langfelder P, Horvath S. WGCNA: an R package for weighted correlation network analysis. BMC bioinformatics. 2008 Dec 29;9(1):559.

Letham DS. Regulators of cell division in plant tissues: V. A comparison of the activities of zeatin and other cytokinins in five bioassays. Planta. 1967 Sep;74(3):228–42.

Li M, Ruwe H, Melzer M, Junker A, Hensel G, Tschiersch H, Schwenkert S, Chamas S, Schmitz-Linneweber C, Börner T, Stein N. The Arabidopsis AAC proteins CIL and CIA2 are sub-functionalized paralogs involved in chloroplast development. Frontiers in Plant Science. 2021 Jun 7;12:681375.

Liebsch D, Juvany M, Li Z, Wang HL, Ziolkowska A, Chrobok D, Boussardon C, Wen X, Law SR, Janečková H, Brouwer B. Metabolic control of arginine and ornithine levels paces the progression of leaf senescence. Plant physiology. 2022 Aug 1;189(4):1943–60.

Lim PO, Kim HJ, Gil Nam H. Leaf senescence. Annu. Rev. Plant Biol.. 2007 Jun 2;58:115–36.

Lobo FD, De Barros MP, Dalmagro HJ, Dalmolin ÂC, Pereira WE, De Souza EC, Vourlitis GL, Rodríguez Ortíz CE. Fitting net photosynthetic light-response curves with Microsoft Excel—a critical look at the models. Photosynthetica. 2013 Sep;51(3):445–56.

Love MI, Huber W, Anders S. Moderated estimation of fold change and dispersion for RNA-seq data with DESeq2. Genome biology. 2014 Dec 5;15(12):550.

Mahapatra K, Mukherjee A, Suyal S, Dar MA, Bhagavatula L, Datta S. Regulation of chloroplast biogenesis, development, and signaling by endogenous and exogenous cues. Physiology and Molecular Biology of Plants. 2024 Feb;30(2):167–83.

Paluch-Lubawa E, Stolarska E, Sobieszczuk-Nowicka E. Dark-induced barley leaf senescence–a crop system for studying senescence and autophagy mechanisms. Frontiers in Plant Science. 2021 Mar 15;12:635619.

Pankasem N, Hsu PK, Lopez BN, Franks PJ, Schroeder JI. Warming triggers stomatal opening by enhancement of photosynthesis and ensuing guard cell CO2 sensing, whereas higher temperatures induce a photosynthesis-uncoupled response. New Phytologist. 2024 Dec;244(5):1847–63.

Pertea M, Kim D, Pertea GM, Leek JT, Salzberg SL. Transcript-level expression analysis of RNA-seq experiments with HISAT, StringTie and Ballgown. Nature protocols. 2016 Sep;11(9):1650–67.

Ritchie ME, Phipson B, Wu DI, Hu Y, Law CW, Shi W, Smyth GK. limma powers differential expression analyses for RNA-sequencing and microarray studies. Nucleic acids research. 2015 Apr 20;43(7):e47-.

Rohart F, Gautier B, Singh A, Lê Cao KA. mixOmics: An R package for ‘omics feature selection and multiple data integration. PLoS computational biology. 2017 Nov 3;13(11):e1005752.

Schippers JH, Schmidt R, Wagstaff C, Jing HC. Living to die and dying to live: the survival strategy behind leaf senescence. Plant physiology. 2015 Oct 1;169(2):914–30.

Sobieszczuk-Nowicka E, Kubala S, Zmienko A, Małecka A, Legocka J. From accumulation to degradation: reprogramming polyamine metabolism facilitates dark-induced senescence in barley leaf cells. Frontiers in plant science. 2016 Jan 6;6:1198.

Sobieszczuk-Nowicka E, Zmienko A, Samelak-Czajka A, Łuczak M, Pietrowska-Borek M, Iorio R, Del Duca S, Figlerowicz M, Legocka J. Dark-induced senescence of barley leaves involves activation of plastid transglutaminases. Amino Acids. 2015 Apr;47(4):825–38.

Steiner E, Israeli A, Gupta R, Shwartz I, Nir I, Leibman-Markus M, Tal L, Farber M, Amsalem Z, Ori N, Müller B. Characterization of the cytokinin sensor TCSv2 in arabidopsis and tomato. Plant Methods. 2020 Nov 16;16(1):152.

Sun CW, Chen LJ, Lin LC, Li HM. Leaf-specific upregulation of chloroplast translocon genes by a CCT motif–containing protein, CIA 2. The Plant Cell. 2001 Sep 1;13(9):2053–61.

Sun CW, Huang YC, Chang HY. CIA2 coordinately up-regulates protein import and synthesis in leaf chloroplasts. Plant Physiology. 2009 Jun 1;150(2):879–88.

Talla SK, Panigrahy M, Kappara S, Nirosha P, Neelamraju S, Ramanan R. Cytokinin delays dark-induced senescence in rice by maintaining the chlorophyll cycle and photosynthetic complexes. Journal of Experimental Botany. 2016 Mar 1;67(6):1839–51.

Thennakoon M, Khanna RR, Singh S, Rashotte AM. cis-Zeatin is an Active Cytokinin that Mediates Gene Expression and Delays Leaf Senescence in Tomato Under Salt Stress. Journal of Plant Growth Regulation. 2026 May;45(5):3686–707.

Tu X, Ren S, Shen W, Li J, Li Y, Li C, Li Y, Zong Z, Xie W, Grierson D, Fei Z. Limited conservation in cross-species comparison of GLK transcription factor binding suggested wide-spread cistrome divergence. Nature Communications. 2022 Dec 9;13(1):7632.

Wang L, Wang Y, Feil R, MacNeill GJ, Lunn JE, Tetlow IJ, Emes MJ. Plastidial starch phosphorylase regulates maltodextrin turnover during starch granule initiation in Arabidopsis leaves. Plant Physiology. 2025 Jun;198(2):kiaf216.

Waters MT, Wang P, Korkaric M, Capper RG, Saunders NJ, Langdale JA. GLK transcription factors coordinate expression of the photosynthetic apparatus in Arabidopsis. The Plant Cell. 2009 Apr 1;21(4):1109–28.

Woo HR, Kim HJ, Lim PO, Nam HG. Leaf senescence: systems and dynamics aspects. Annual review of plant biology. 2019 Apr 29;70(1):347–76.

Xie M, Chen H, Huang L, O’Neil RC, Shokhirev MN, Ecker JR. A B-ARR-mediated cytokinin transcriptional network directs hormone cross-regulation and shoot development. Nature communications. 2018 Apr 23;9(1):1604.

Xing J, Pan J, Yang W. Chloroplast protein translocation complexes and their regulation. Journal of Integrative Plant Biology. 2025 Apr;67(4):912–25.

Yang CY, Sun CW. Sequence analysis and protein interactions of Arabidopsis CIA2 and CIL proteins. Botanical studies. 2020 Jun 18;61(1):20.

Yang CY, Yan WY, Chang HY, Sun CW. Arabidopsis CIA2 and CIL have distinct and overlapping functions in regulating chloroplast and flower development. Plant Direct. 2022 Jan;6(1):e380.

Zarrin Ghalami R, Burdiak P, Kamran M, Duszyn M, Rusaczonek A, Muszyńska E, Karpiński S. Role of CIA2 and CIL in the Regulation of Chloroplast Development During Photomorphogenesis in Arabidopsis. Cells. 2026 Feb 11;15(4):333.

Zhang Z, Li W, Gao X, Xu M, Guo Y. DEAR4, a member of DREB/CBF family, positively regulates leaf senescence and response to multiple stressors in Arabidopsis thaliana. Frontiers in Plant Science. 2020 Mar 31;11:367.

Zhao Y, Zhang Y, Li S, Tan S, Cao J, Wang HL, Luo J, Guo H, Zhang Z, Li Z. Leaf senescence database v5. 0: a comprehensive repository for facilitating plant senescence research. Journal of Molecular Biology. 2024 Sep 1;436(17):168530.

Zubo YO, Blakley IC, Yamburenko MV, Worthen JM, Street IH, Franco-Zorrilla JM, Zhang W, Hill K, Raines T, Solano R, Kieber JJ. Cytokinin induces genome-wide binding of the type-B response regulator ARR10 to regulate growth and development in Arabidopsis. Proceedings of the National Academy of Sciences. 2017 Jul 18;114(29):E5995–6004.

Zwack PJ, Rashotte AM. Cytokinin inhibition of leaf senescence. Plant signaling & behavior. 2013 Jul 1;8(7):e24737.

