## Supplemental Figures for "Cytokinin Mediates Photosynthesis and Senescence Action Through CHLOROPLAST IMPORT APPARATUS 2 (CIA2) and CIA2-like (CIL)"

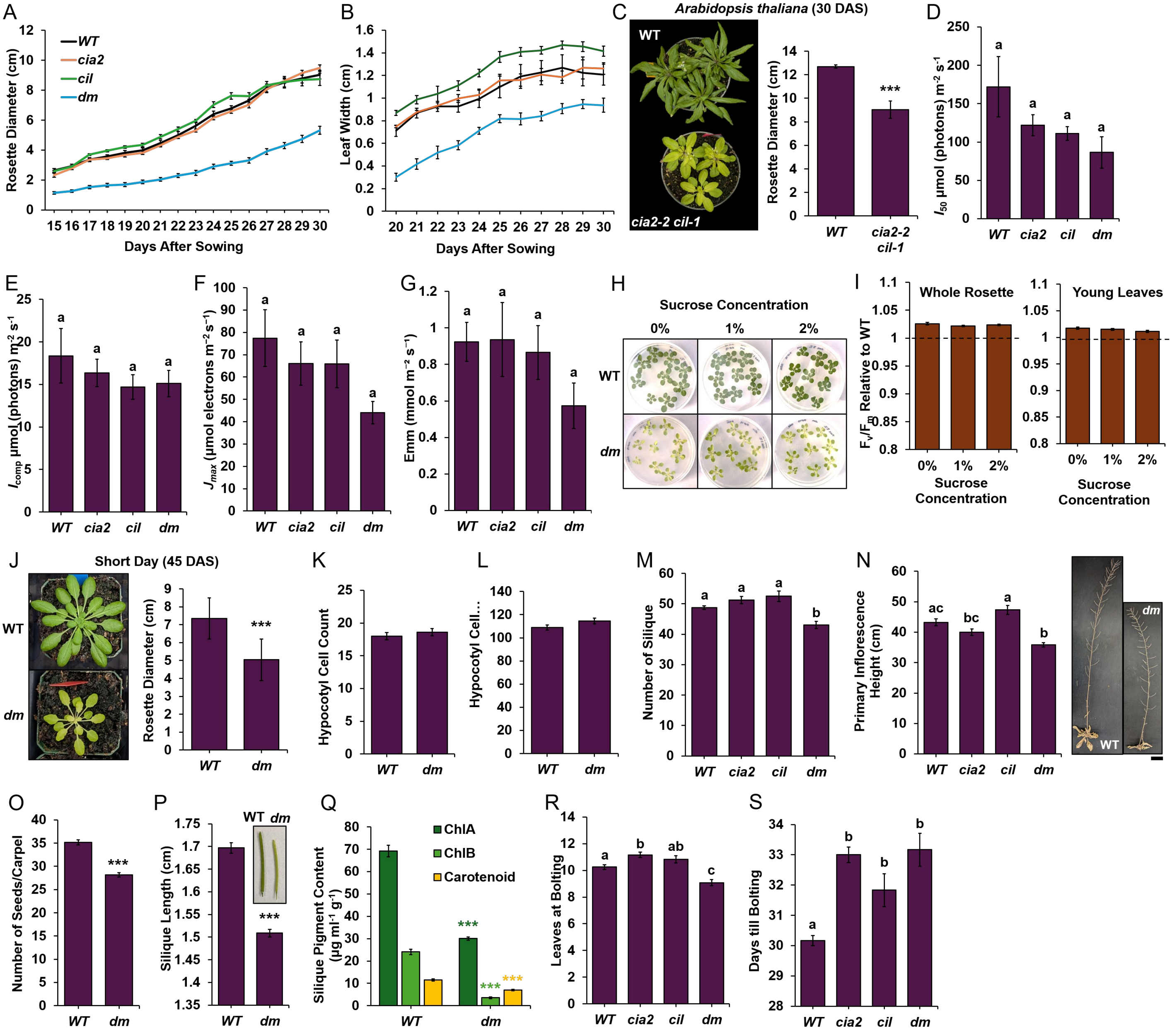


**Supp Fig 1:** Phenotypic characterization and photosynthesis analysis of Arabidopsis genotypes. (A) Using image-based phenotyping, rosette diameter was recorded 15-30 days after sowing (DAS) while (B) leaf width noted from 20-30 DAS (n = 6). (C) WT and *cia2-2 cil-1* (T-DNA insertion mutants) were measured for rosette diameter (30 DAS), using 6 plants per genotype. (D-E) $I_{50}$ and $I_{compensation}$ (µmol photons $m^{-2}$ $s^{-1}$) performed based on light-curve analysis. (F) Rate of electron transport ($J_{max}$; µmol electrons $m^{-2}s^{-1}$) measured using $AC_i$ curve, normalizing values to 25 °C before presenting. (G) Transpiration rate (Emm; mmol $m^{-2}$ $s^{-1}$) recorded at 1855 µmol $m^{-2}$ $s^{-1}$ PAR. Leaf 5-8 were utilized for all LI6800-based photosynthesis analysis using at least 6 plants at 45 DAS. (H) Seedlings 5 DAG were moved to MS media with 0%, 1%, or 2% sucrose for 7 days following which they were photographed and (I) $F_v/F_m$ recorded for whole rosette or young leaves (n = 24). Dashed line representing WT $F_v/F_m$ levels. (J) WT and *dm* were grown in short day (8:16 h light:dark) and 45 DAS the rosette diameter was recorded using ImageJ analysis, measuring 9 plants per genotype. Representative plants shown. (K-L) Hypocotyl epidermal cells were observed using 7 days after germination (DAG) seedlings. ImageJ was used to perform cell counts and cell lengths across 12 samples per genotypes. Cell lengths of 216 and 223 cells were measured in WT and *dm* respectively. (M-N) Number of siliques on primary inflorescence and primary inflorescence height (in cm) at 80 DAS (n = 8). Representative pictures after excising secondary inflorescences. Scale Bar = 3 cm. (O-P) Number of seeds per carpel and silique length noted for silique 5 – 10 (from the bottom) at 45 DAS. Representative siliques shown. (Q) Silique pigment content at 45 DAS, pooling 10 siliques per sample (n = 6). (R) Total number of leaves on the day of bolting (n = 12). (S) Number of days until bolting starting from DAS (n = 6). All results are presented as mean ±SE performed across three biological replicates. Statistical tests performed using ANOVA and Tukey's HSD with different letters noting statistically significant difference (p.adj < 0.05). Student's t test performed for (C, D, O, P, and Q) with asterisk (*** p < 0.005).

A

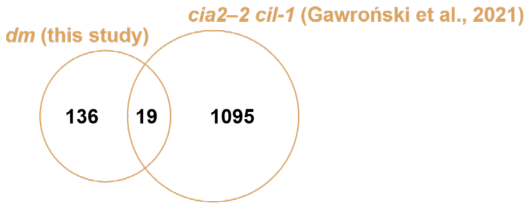


B

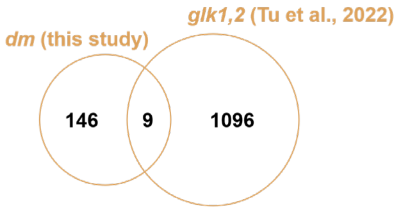


**Table: List of 19 shared DEGs between this study and Gawroński et al., 2021**

| Gene ID | Symbol | Description |
|---|---|---|
| AT5G02500 | HSC70-1 | heat shock cognate protein 70-1 |
| AT1G06540 | | - |
| AT2G12190 | | Cytochrome P450 superfamily protein |
| AT4G13560 | UNE15 | Late embryogenesis abundant protein (LEA) family protein |
| AT4G13570 | HTA4 | histone H2A 4 |
| AT4G13572 | | - |
| AT4G13575 | | - |
| AT4G13577 | | hypothetical protein |
| AT2G20570 | GLK1/GPRI1 | GBF's pro-rich region-interacting factor 1 |
| AT4G20140 | GSO1 | Leucine-rich repeat transmembrane protein kinase |
| AT5G38420 | RBCS2B | Ribulose bisphosphate carboxylase (small chain) family protein |
| AT3G55240 | | Plant protein 1589 of unknown function |
| AT3G55290 | | NAD(P)-binding Rossmann-fold superfamily protein |
| AT5G52970 | | thylakoid lumen 15.0 kDa protein |
| AT3G58430 | | TRAF-like family protein |
| AT5G55930 | OPT1 | oligopeptide transporter 1 |
| AT5G58310 | MES18 | methyl esterase 18 |
| AT1G75230 | | DNA glycosylase superfamily protein |
| AT1G75945 | | hypothetical protein |

**Table: List of 9 shared DEGs between *dm* (this study) and *glk1,2* (Tu et al., 2022)**

| Gene ID | Symbol | Description |
|---|---|---|
| AT5G01600 | FER1 | ferretin 1 |
| AT1G12310 | | Calcium-binding EF-hand family protein |
| AT4G13570 | HTA4 | histone H2A 4 |
| AT4G13572 | | None |
| AT4G13575 | | None |
| AT2G20570 | GLK1/GPRI1 | GBF's pro-rich region-interacting factor 1 |
| AT5G38420 | RBCS2B | Ribulose bisphosphate carboxylase (small chain) family protein |
| AT3G51860 | CAX3 | cation exchanger 3 |
| AT3G55240 | | Plant protein 1589 of unknown function |

**Supp Fig 2:** (A) Differentially expressed genes (DEGs) from *dm* vs WT comparisons noted for this study (using *cia2-1 cil-5*; mature leaf 5-6 on day of bolting) and Gawroński et al., 2021 (using *cia2–2 cil-1*; young leaves) along with list of 19 shared genes. (B) DEGs from *dm* vs WT comparison (this study) and *glk1,2* vs WT comparison from Tu et al., 2022 along with list of 9 shared genes.

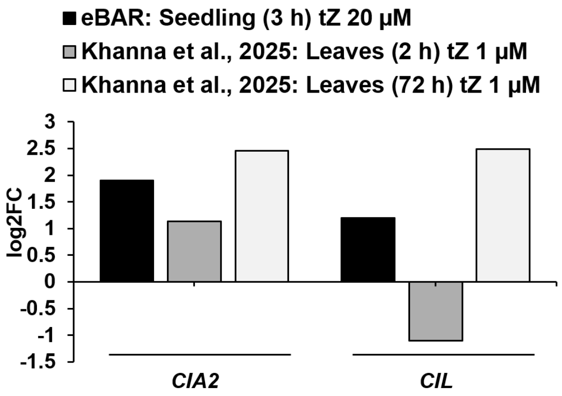


**Supp Fig 3:** eBAR seedling (3 h) *trans*-Zeatin 20 µM and Khanna et al., 2025 leaves (2 h and 72 h) *tZ* 1 µM was used to evaluate *CIA2* and *CIL* expression, presented on a log2FC scale.

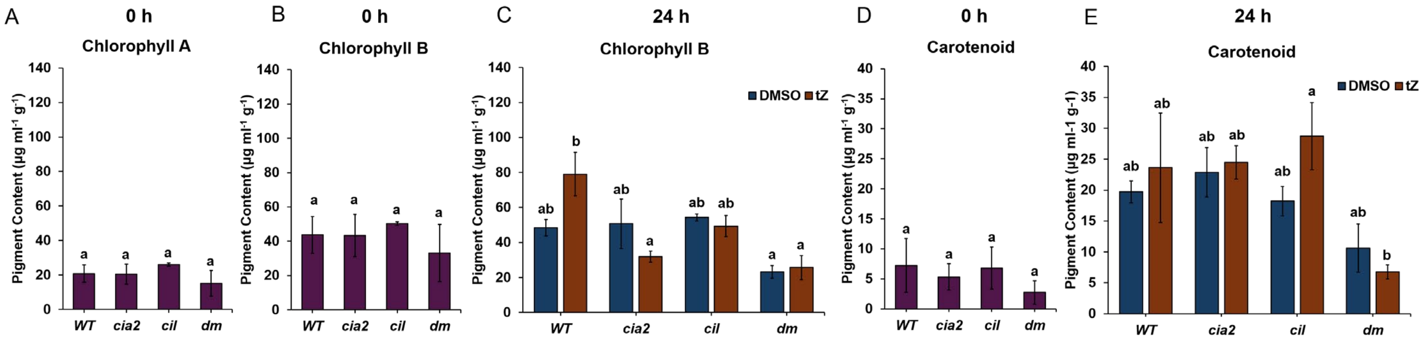


**Supp Fig 4:** (A-E) Chlorophyll a (A), chlorophyll b (B-C), and carotenoid (D-E) content were calculated using dark-grown seedlings (5 DAG) that were light-exposed for 0 h or 24 h under *tZ* (1 µM) treated or DMSO conditions (noted using colors). Pigment was extracted using 30 whole-seedlings pooled for each sample performing at least three biological replicates. All results are represented as mean ±SE. Statistical test performed using ANOVA and Tukey's HSD with different letters noting significant difference (p.adj < 0.05).

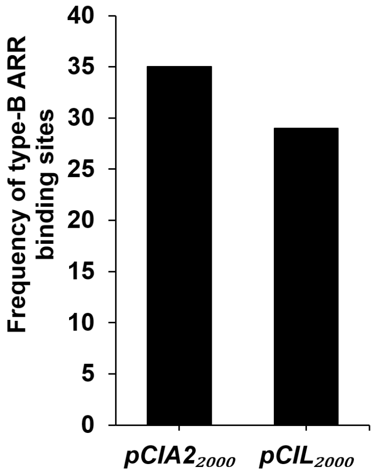


**Supp Fig 5:** Frequency of type-B ARR (RRBs) binding site in promoter elements of *CIA2* and *CIL* (2000 bp upstream from start).

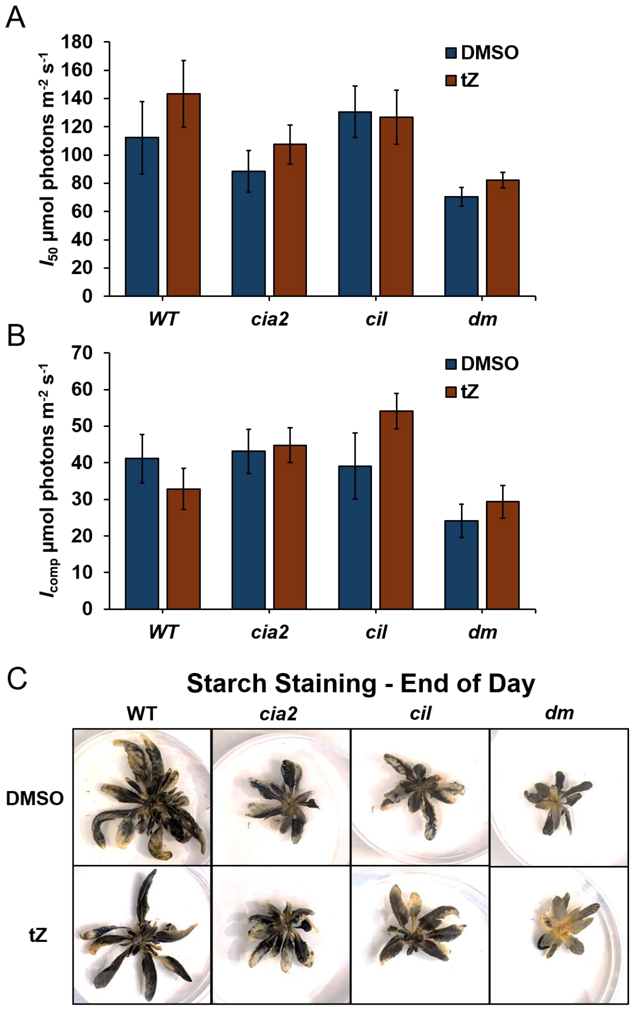


**Supp Fig 6:** (A-B) $I_{50}$ and $I_{compensation}$ (µmol photons m$^{-2}$ s$^{-1}$) performed based on light-curve analysis at 60 DAS following *tZ* (µM) watering. No statistically significant difference was observed between *tZ* vs DMSO treatments. Leaf 5-8 were utilized for all LI6800-based photosynthesis analysis using at least 5 plants measured on 60 DAS across three biological replicates. (C) Starch accumulation staining rosettes using Lugol's stain at end of day (30 DAS). Representative photographs shown.

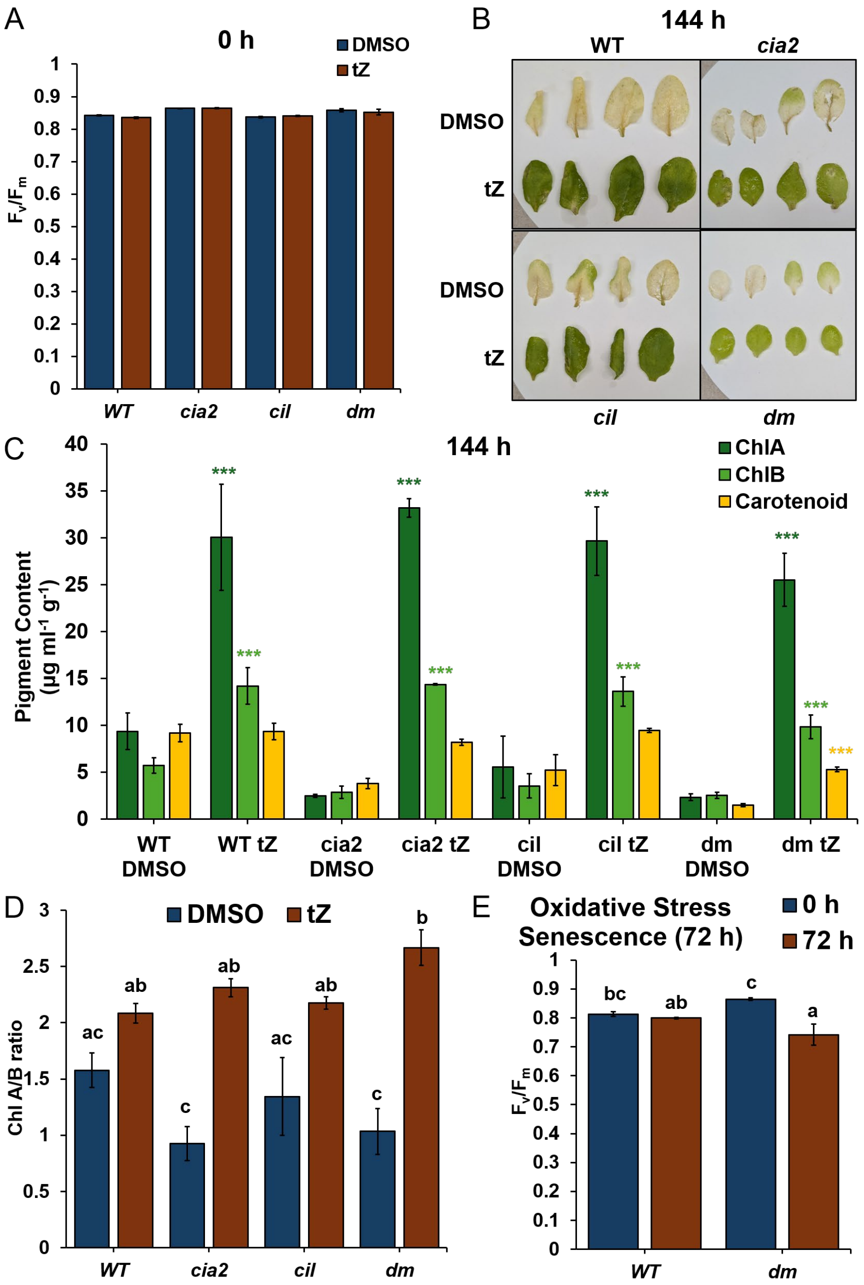


**Supp Fig 7:** (A) Photosystem II efficiency ($F_v/F_m$) analysis using leaves 3-8 under dark-induced senescence (DIS) with *tZ* (1 µM) treatment or DMSO conditions (noted using colors) at 0 h. 21-31 leaves were measured per genotype and treatment. (B) Leaves 3-6 following DIS (144 h) under *tZ* (1 µM) treatment or DMSO conditions. Representative leaves shown. (C-D) Pigment content analysis using leaves 3-4 under dark-induced senescence (DIS) 144 h with *tZ* (1 µM) DMSO, with (C) showing chlorophyll a, chlorophyll b, and carotenoid content (noted using colors) and (D) showing chlorophyll a/b ratio. 3-9 leaves were measured per genotype and treatment. (E) WT or *dm* leaf 5-6 treated with $H_2O_2$ (20 mM) were recorded for $F_v/F_m$ at 0 h and 72 h (noted with color), measuring at least 5 leaves for each genotype per treatment. All results represented as mean ±SE for at least three biological replicates. Statistical test performed using ANOVA and Tukey's HSD with different letters noting significant difference (p.adj < 0.05) or with asterisk (*** p < 0.005).

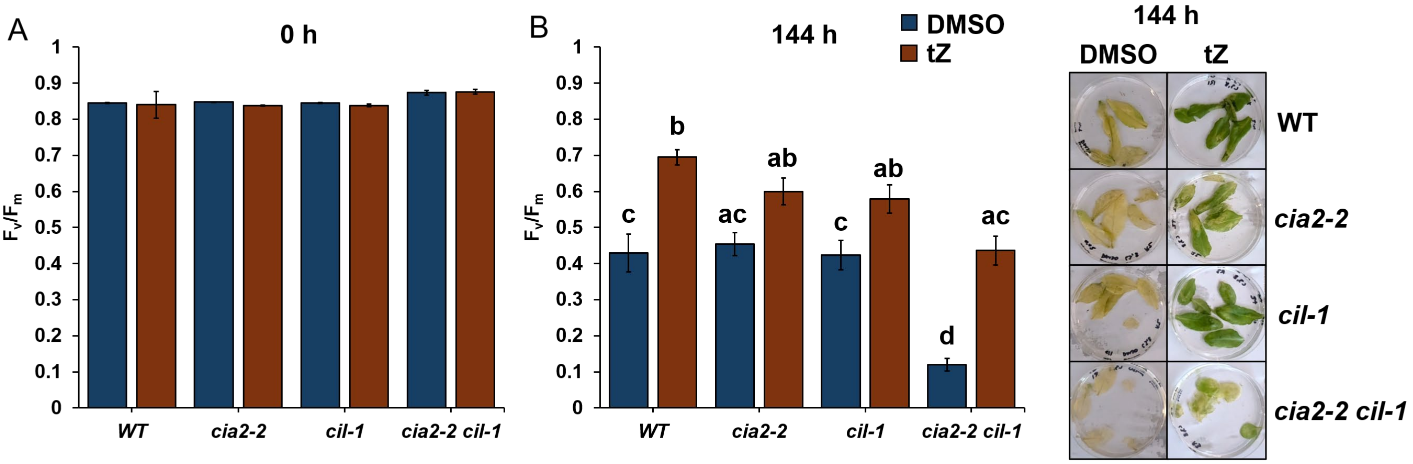


**Supp Fig 8:** (A-B) Photosystem II efficiency ($F_v/F_m$) analysis using leaves 5-6 under dark-induced senescence (DIS) with *tZ* (1 µM) or DMSO (noted using colors) at 0 h (A) or 144 h (B). 15-47 leaves were measured per genotype and treatment. All results represented as mean ±SE for at least three biological replicates. Statistical test performed using ANOVA and Tukey's HSD with different letters noting statistically significant difference (p.adj < 0.05). Representative leaves are shown.

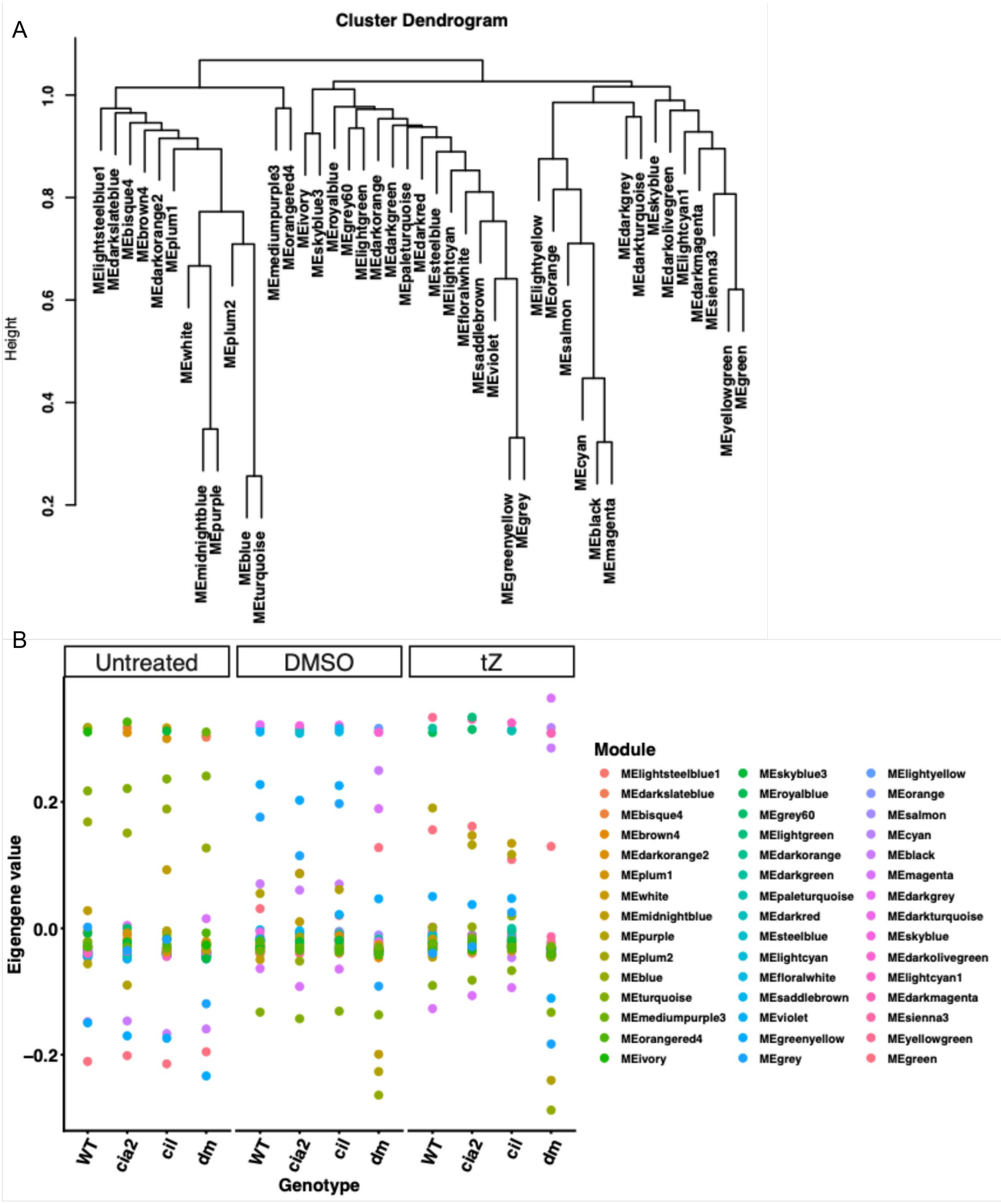


**Supp Fig 9:** (A) Hierarchical clustering of 45 WGCNA gene modules each named with a unique color. (B) WGCNA gene modules and their eigengene values across genotypes plotted for each treatment – 0 h (untreated) and 72 h DIS with *tZ* or DMSO. Each module is presented as a dot with a color as noted in legend.

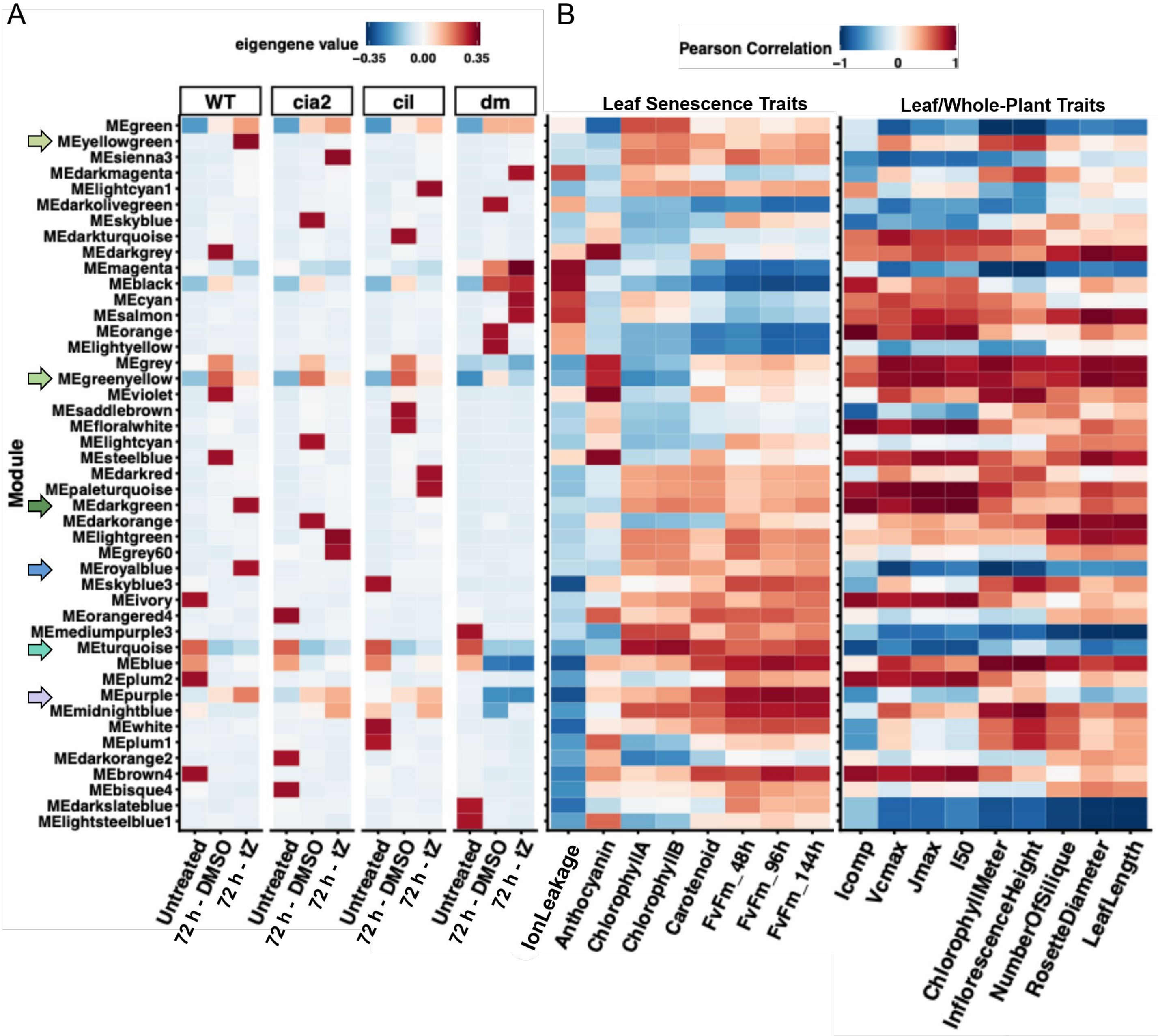


**Supp Fig 10:** (A) Heatmap representing WGCNA modules (each named with a unique color) eigengene expression (noted with red/blue color gradient) across genotypes in untreated (0 h) and DIS 72 h *tZ* (1 μM) treatment or DMSO conditions. Module arranged according to hierarchical clustering and those showing expression changes in a cytokinin-specific manner (in WT but lacking in *dm*) are pointed out using colored arrows. (B) Correlation analysis performed on WGCNA module eigengene expression and physiological traits under DIS (leaf senescence traits) or growth/photosynthesis (leaf or whole-plant traits). Traits clustered on X-axis based on similarity in correlation values. Red and blue colors of the heatmap depict positive or negative correlation (Pearson correlation ranging from 1 to -1).

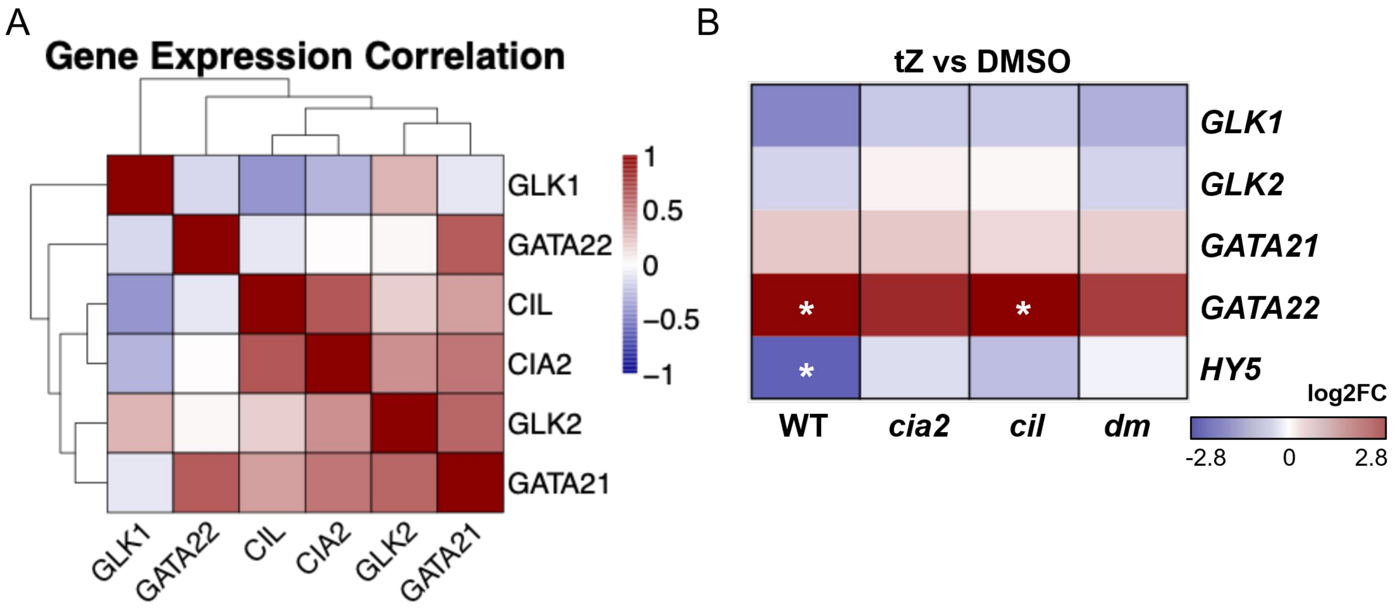


**Supp Fig 11:** (A) Correlation analysis of *CIA2, CIL, GLK1, GLK2, GATA21,* and *GATA22* expression patterns, using gene counts across all genotypes and treatments. Red and blue colors of the heatmap depict positive or negative correlation (Pearson correlation ranging from 1 to -1). (B) Heatmap representing expression profile of selective genes on a log2FC scale (*t*Z vs DMSO comparison) with red depicting induction while blue depicting repression. Asterisk used to note statistically significant difference (FDR < 0.05).

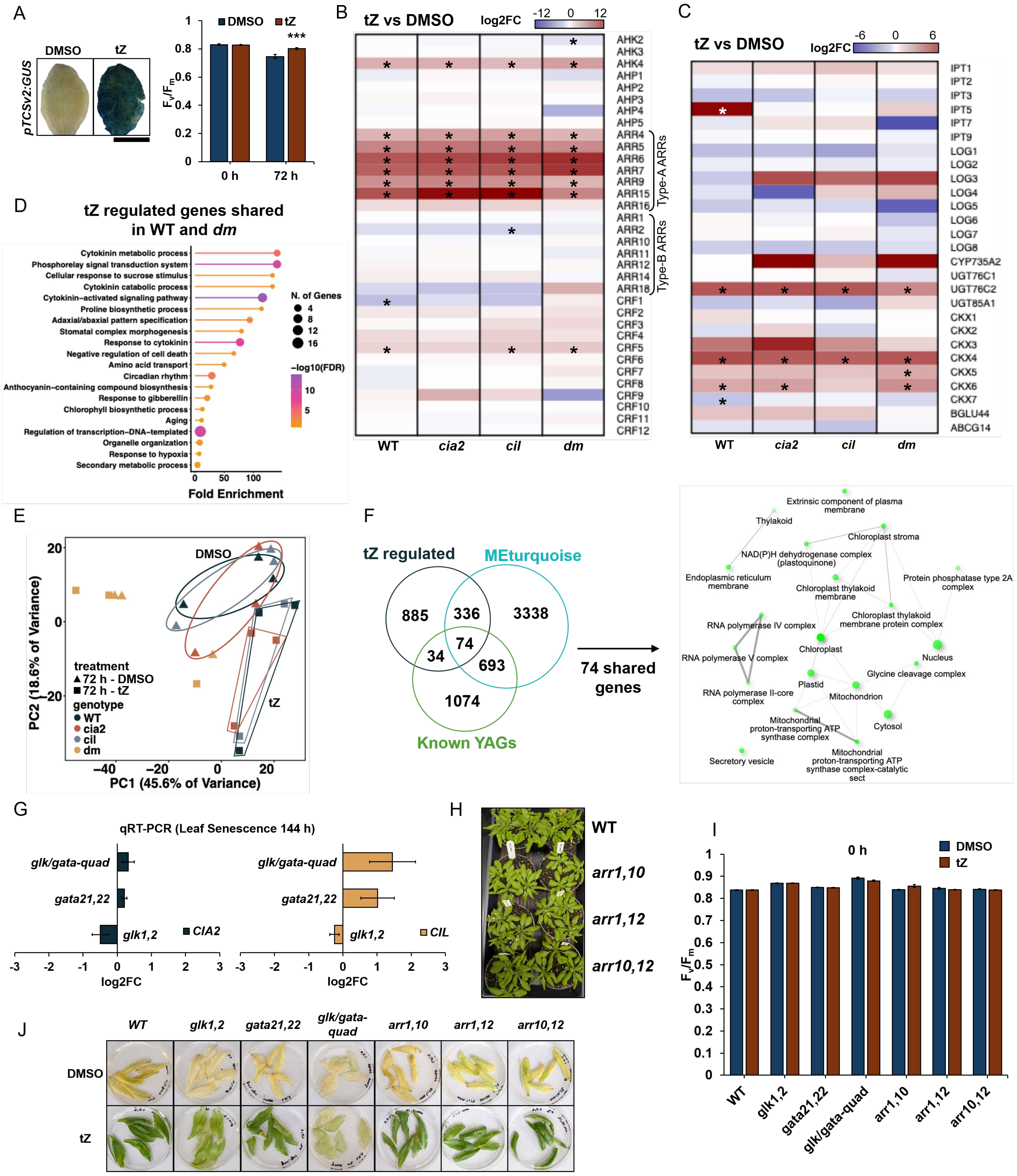


**Supp Fig 12:** (A) Leaves 5-6 from *pTCSv2:GUS* following DIS (72 h) with *tZ* (1 µM) or DMSO were GUS stained. Representative leaves shown. Scale bar = 1 cm. *pTCSv2:GUS* leaves were recorded for $F_v/F_m$ levels at 0 h and 72 h (colors noting *tZ* or DMSO), measuring at least 9 leaves per treatment. Data presented as mean ±SE across three biological replicates. Statistical test performed using Student's t test with asterisk (*** $p < 0.005$) noting significant difference. (B-C) Heatmap represents expression profile of cytokinin-signaling (B) and cytokinin-metabolism (C) genes. Gene expression represented on a log2FC scale (*tZ* vs DMSO comparison) with red depicting induction while blue depicting repression. Asterisk used to note statistically significant difference (FDR < 0.05). (D) Lollipop plot representing fold enrichment for 89 genes commonly regulated by *tZ* in WT and *dm*. Selective GO terms presented with dot size used to note number of genes within each GO term and color noting -log(FDR). (E) PCA depicting overall gene expression differences at 72 h under DIS with *tZ* or DMSO. Each dot represents a single sample with genotype and treatment noted with color and shape, respectively. Oval and quadrilateral denote DMSO and *tZ* treatment grouping respectively with color of outline depicting a distinct genotype: WT, *cia2*, *cil*. (F) Venn diagram denoting *tZ* regulated genes in WT, MEturquoise, and known youth-associated genes (YAGs) from Guo et al., 2025. Shared genes amongst these were used to evaluate GO terms (cellular components), with size of node denoting number of genes and branches noting shared genes across GO category. (G) qRT-PCR analysis performed for leaf 5-6 under DIS (144 h). Leaves 5-6 were pooled from three plants (total 6 leaves) for each replicate and flash frozen. Data presented on log2FC scale as mean ±SE relative to WT across three biological replicates, with colors noting *CIA2* or *CIL* gene expression. (H) Genotypes were grown and photographed on 30 DAS. Representative plants shown. (I) PSII efficiency ($F_v/F_m$) analysis measuring leaves 5-6 at DIS 0 h with *tZ* or DMSO (noted using colors). 17-22 leaves were measured (at 72 h 4-6 leaves measured) with results represented as mean ±SE across three biological replicates. (J) Leaves 5-6 across genotypes at 144 h (DIS) with *tZ* or DMSO. Representative pictures shown.

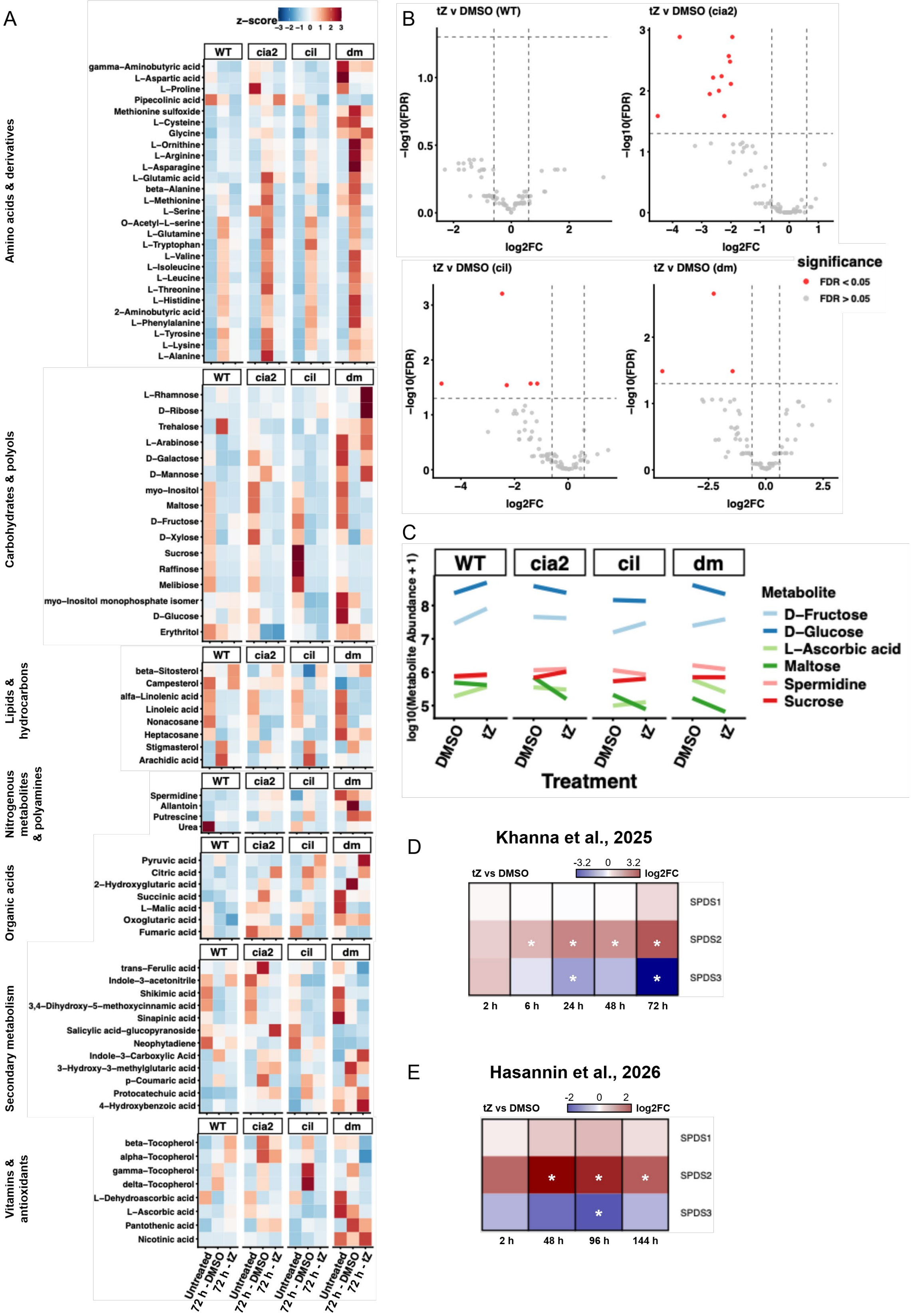


**Supp Fig 13:** (A) Heatmaps depicting normalized metabolite abundance (presented on a z-score scale) averaged of four biological replicates across genotypes under DIS 0 h (untreated) and 72 h treated with *tZ* or DMSO. Metabolites on Y-axis arranged according to z-score based hierarchical clustering. Red and blue indicate metabolite induction and repression respectively. (B) Volcano plots used to note change in metabolite abundance (on a log2FC scale) under *tZ* vs DMSO comparison. Each point denotes a distinct metabolite with differentially abundant metabolites (DAMs; FDR < 0.05) noted in red; indicated above the -log10(FDR) dashed horizontal line scale. (C) Line graphs showing average metabolite abundance on a log10(metabolite abundance +1) scale of selective metabolites that show a cytokinin-response in WT but lack similar trends in *dm*. Data presented as average of four biological replicates, with each metabolite shown with a unique color (as noted in legend). (D-E) Heatmap representing expression profile of spermidine-metabolism genes, presented from Khanna et al., 2025 (D) and Hasannin et al., 2026 (E) on a log2FC scale (*tZ* vs DMSO comparison) with red depicting induction while blue depicting repression. Asterisk used to note significant difference (FDR < 0.05).

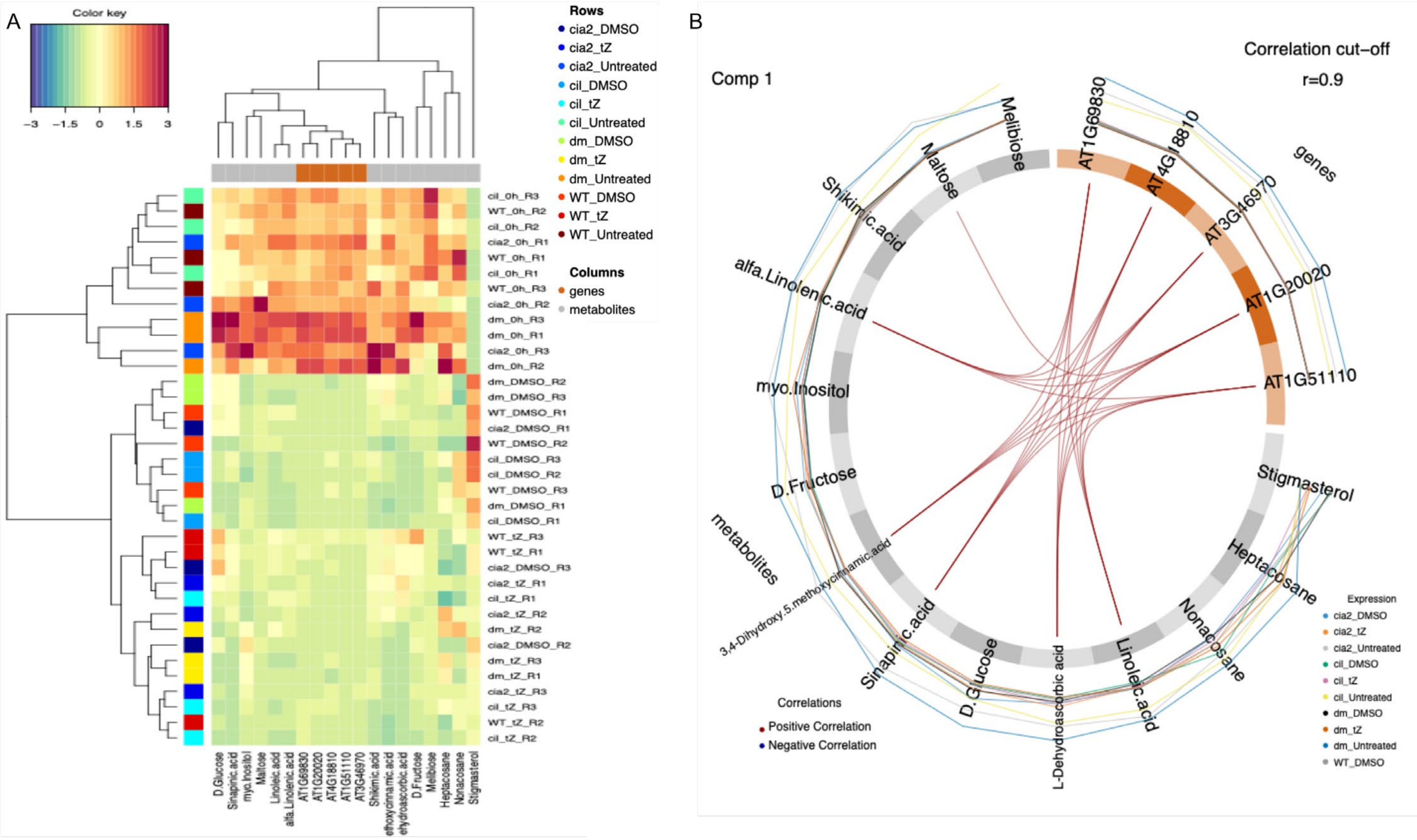


**Supp Fig 14:** Integrated transcriptomics-metabolomics analysis performed using mixOmics package. A multiblock sparse partial least squares discriminant analysis (sPLS-DA) was used to integrate transcripts per million and normalized metabolite abundance, generating a Clustered Image Map (CIM; A) and circus plot (B) for component 1, with genes-metabolite correlation cutoff = 0.9. In the CIM plot, the genes/metabolites (X-axis) and biological sample (Y-axis) were clustered and depicted using color as noted in figure legend.

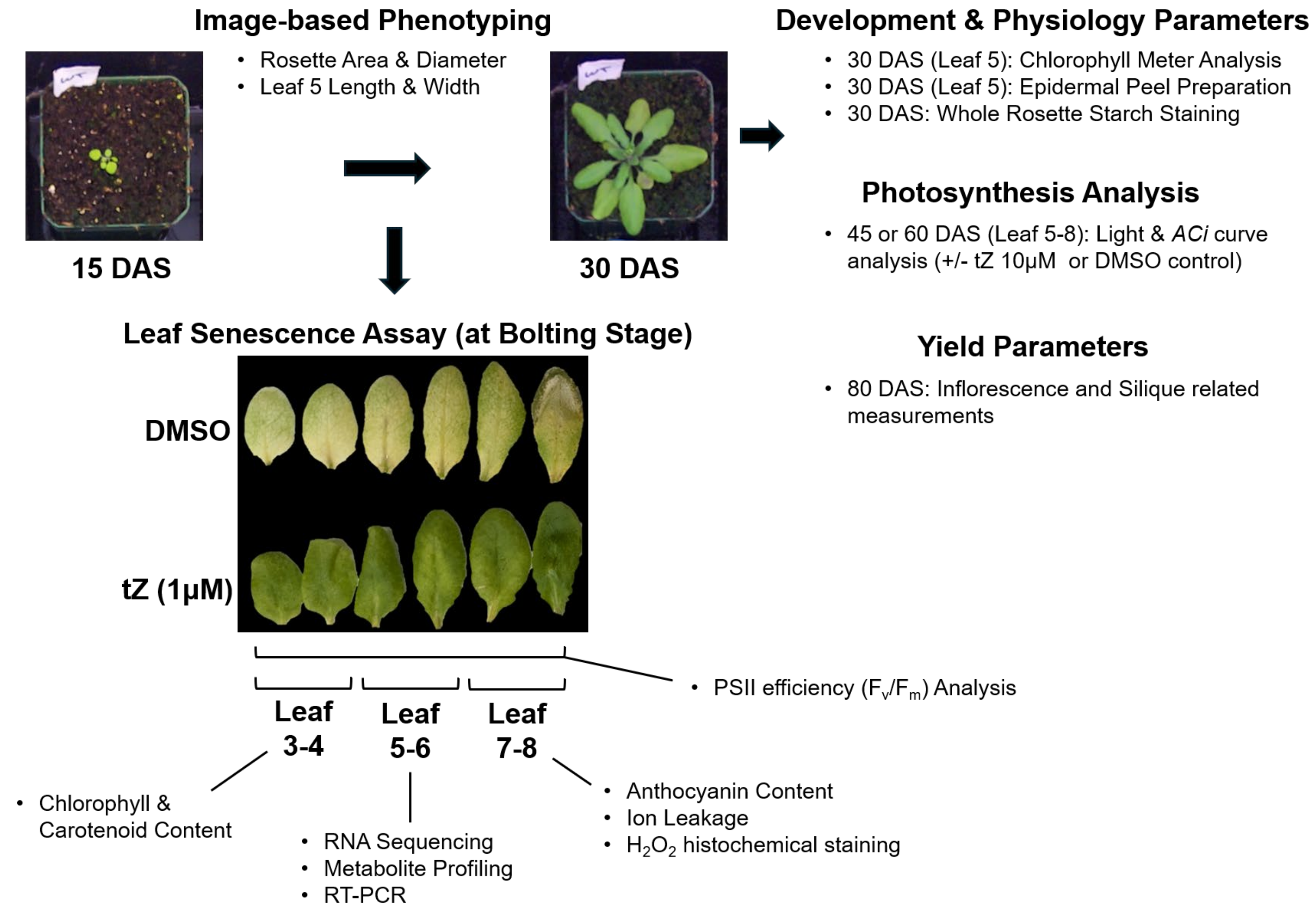


**Supp Fig 15: Methodology utilized for whole-plant and senescence analysis.** This indicates the phenotyping plan followed for image-based phenotyping, growth, yield-related parameters, physiological testing, transcriptomics, and metabolomics approaches. Senescence phenotyping modified from Bresson et al., 2018.
