## Supplemental Methods for "Cytokinin Mediates Photosynthesis and Senescence Action Through CHLOROPLAST IMPORT APPARATUS 2 (CIA2) and CIA2-like (CIL)"

**Supplementary methods**

*Image based phenotyping and growth-related parameters*

Raspberry Pi cameras were utilized to image Arabidopsis rosettes from 15 – 30 days after sowing (DAS). Rosette and leaf growth parameters were evaluated using ImageJ software. For Rosette diameter the longest diameter between two leaf tips was utilized. For leaf length, the distance from leaf tip to base was measured while for leaf width the distance across the broadest portion of the leaf was measured. Rosette size was analyzed by thresholding function or manually tracing the entire rosette and calculating covered area. Handheld chlorophyll meter (atLeaf, DE, USA) was utilized for non-destructive analysis of pigment content – measuring leaf 5 from 30 DAS plants.

For bolting analysis, seeds were kept in 4 °C for 72 h to stratify before sowing on planting mix. Number of days from sowing to bolting and number of leaves on the day of bolting were recorded. For yield-related parameters, rosettes were detached from 80 DAS plants and all secondary inflorescences excised. Plants were photographed on a black background and length of primary inflorescence and silique size recorded using ImageJ software. For silique size, silique 5 to 10 from the bottom were used. Further, number of siliques on primary inflorescence were counted. Siliques were ripped open and number of seeds per carpel were counted under a Nikon SMZ1500 stereomicroscope.

*Cell size and stomata parameters*

Leaf 5 was excised from 30 DAS soil-grown plants. Epidermal peels (using clear nail polish) were visualized under Nikon Eclipse 80i microscope at 20x objective magnification. Images were captured at three different positions including leaf tip, middle, and base. Epidermal cell size and stomata lengths were measured using ImageJ software. Number of cells across field width and number of stomata per field were recorded.

For analysis of hypocotyl cells, seeds were surface sterilized and kept in 4 °C for 72 h. Seeds were sown on MS 1x with Sucrose (1%) and kept in 22 °C /18 °C for 16 h/8 h day/night cycle. 5 DAS seedlings were moved to be vertically oriented and grown for 7 more days after which hypocotyl cells were observed under Nikon Eclipse 80i microscope (10x objective magnification). Cell length measurements were performed using ImageJ software, measuring lengths of cell position 5-10.

*Leaf senescence assay and physiology-related parameters*

On the day of bolting, leaves were excised and utilized for dark-induced leaf senescence (DIS) assay as per Johnston et al., 2026. Briefly, specific leaf numbers (Fig 9; or in figure legends) were floated on MES buffer 3 mM pH 5.7 supplemented with *trans*-Zeatin (*t*Z; 1 μM) or DMSO control. *t*Z was obtained from PlantMedia (USA) with greater than 90% purity. Plates were wrapped with aluminum foil to maintain dark conditions and returned to 22 °C /18 °C plant growth chamber conditions. Leaves were non-destructively sampled for F_v_/F_m_ or pigment/oxidative stress parameters as described below.

For photosystem II efficiency analysis, leaves were dark adapted for at least 30 min following which F_v_/F_m_ measurements were recorded at various timepoints (as described in text) using Open FluorCam (Photon Systems Instruments). For pigment analysis, after 144 h of DIS - Leaf 3 and 4 were pooled and weighted together. Leaves were then transferred to pure methanol (5 ml) and kept in 4 °C overnight. Using 1 ml of pigment extract, A_470_, A_652_, and A_665_ were recorded with Beckman DU730 UV/Vis spectrophotometer. Pigment content including chlorophyll A, chlorophyll B, and carotenoids was calculated using a previously described formula (Sumanta et al., 2014).

For anthocyanin content analysis leaf 7 and 8 after 144 h of DIS were pooled together and weighed. Leaves were transferred to MeOH:water:HCL (80:20:1) solution (5 ml) and kept in 4 °C overnight. Using 1 ml of pigment extract, A_530_ and A_657_ were recorded. Anthocyanin concentration was calculated as the formula A_530_ - (1/3 * A_657_) as previously described (Lindoo and Caldwell, 1978). For ion leakage analysis, leaves 7 and 8 were pooled and shaken in 10 ml of DI water for 2 h. Using an electrical conductivity meter an initial reading was recorded. Leaves were then boiled for 30 min and final reading recorded. Ion leakage was reported as a percentage = (Initial reading/Final reading) * 100 (Zhang et al., 2020). For H_2_O_2_ histochemical staining the Daudi and O’Brien et al., 2012 protocol was utilized. Briefly, leaves 7 and 8 were transferred to 3,3'-diaminobenzidine (DAB) stain solution and vacuum infiltrated for 30 min followed by overnight incubation. Leaves were cleared using ethanol:acetic acid:glycerol (3:1:1) solution in a boiling water bath (95 °C) for 15 mins. Following replacement of clearing solution, leaves were imaged using a Nikon SMZ1500 stereomicroscope, and photographed with a Qimaging Fast 1394 digital camera.

For RT-PCR analysis, leaves 5 and 6 were flash frozen after 144 h of DIS and stored in -80 °C until grinding was performed. This was followed by RNA-isolation similar to that described under transcriptomic analysis.

*Cytokinin watering and photosynthesis analysis*

Soil-grown plants were tested for photosynthesis analysis at 45 DAS. Alternatively, plants were watered with *t*Z (10 μM) or DMSO dissolved in water applying 50 ml per pot every 5 days starting from 20 DAS until 60 DAS. Photosynthesis and CO_2_ exchange parameters were measured using LI-6800 Portable Photosynthesis System (LICOR biosciences. Lincoln, Nebraska, USA) at 45 DAS or 60 DAS as mentioned in text. The chamber temperature, light, and humidity were set to match the conditions of the plant growth chamber. This included setting temperature to 22 °C and relative humidity to 55%. Air flow was set to 400 μmol s^-1^ and fan speed to 10,000 rpm. Referring to previously described work (Eckstein et al., 2012), for light curve analysis the following light intensities were utilized: 0, 5, 55, 85, 115, 145, 205, 305, 435, 595, 845, 1245, 1645, and 1855 PAR (μmol m^-2^ sec^-1^) – all parameters measured at 425 μmol mol^-1^ CO_2_ concentration. For *A*C*_i_* curves, light was set to 1300 μmol m^-2^ sec^-1^ since it was the point of *A* saturation as noted in the light curves, and modifying from previously described work (Pankasem et al., 2024) the following sequential reference CO_2_ concentrations were used: 425, 300, 200, 100, 50, 25, 425, 600, 800, 1000, 1200, and 1500 μmol mol^-1^. Analysis of net photosynthetic light-response curve was performed as per Lobo et al., 2013, generating *I_50_* and *I_comp_* using the model described in Kaipiainen, 2009. Using *AC_i_* curves, we calculated maximum rate of rubisco carboxylation (*V_cmax_*) and rate of electron transport (*J_max_*) as per the model described in Bernacchi et al., 2001, 2003. Measurements were corrected to 25 °C before plotting.

*Starch staining*

Soil-grown plants that were *t*Z (10 μM) or DMSO watered were sampled at 30 DAS for starch distribution analysis at end of day and end of night periods. Briefly, entire rosettes were detached and added to boiling water for 5 mins – to kill any biological activity. This was followed by transferring rosettes to boiling ethanol for 5 mins removing any pigments and then to Lugol’s stain. Rosettes were destained using distilled water and photographed. Starch staining leaves were performed similarly utilizing leaf 5-6 after 144 h of DIS. Leaves were imaged using a Nikon SMZ1500 stereomicroscope, and photographed with a Qimaging Fast 1394 digital camera. All pictures presenting without altering the original image integrity.

*Sucrose gradient and photoperiod analysis*

Seeds were surface sterilized and kept in 4 °C for three days. After stratification, seeds were plated on MS 1x with 1% sucrose. 5 DAS, they were moved to MS media containing 0%, 1% and 2% sucrose. After 7 days, pictures were taken to perform rosette diameter using ImageJ software and F_v_/F_m_ analysis performed. For photoperiod analysis, stratified seeds on MS media plates were kept in long day conditions. 5 DAS, they were moved to short day (8 h/16 h day/night), long day (16 h/8 h day/night), or continuous light conditions (24 h day) for 7 additional days. Media plates were dark adapted for 30 mins and F_v_/F_m_ recorded.

*De-etiolation assay*

After stratification, seeds on MS media plates were wrapped with aluminum foil to maintain dark conditions and kept in 22 °C/18 °C for 7 days. Etiolated seedlings were pooled (30 seedlings) and weighed followed by pigment extraction marking the 0 h reading. Similarly, seedlings were transferred to 3 mM MES containing *t*Z (1 μM) or DMSO and transferred to light conditions while gently shaking. Following 24 h of incubation, 30 seedlings were utilized for pigment extraction. Similarly, 30 seedlings were flash frozen with liquid nitrogen and stored in -80 °C until grinding and RNA isolation.

*qRT-PCR analysis*

Leaf 5-6 were excised on day of bolting, and flash frozen in liquid nitrogen and stored in -80 °C until grinding. Leaves 5-6 were pooled from three plants (total 6 leaves) for each biological replicate performing a total of three biological replicates. RNA isolation was performed using at least 50 mg of frozen tissue powder utilizing RNeasy Plant Mini Kits (Qiagen) according to manufacturer's instructions. Following quantification using nanodrop spectrophotometer, cDNA was synthesized using qScript cDNA Synthesis Kit (QuantaBio). qRT-PCR analysis was performed using PerfeCTa SYBR® Green FastMix (QuantaBio) and relative expression performed using 2^-ΔΔCT^ method, with all primer sequences utilized in the analysis being mentioned in Supp File 6.

*Promoter expression analysis*

The *pTCSv2:GUS* lines (Steiner et al., 2020) were grown and leaves 5-6 excised to perform DIS. At 72 h, leaves were transferred to GUS staining solution and vacuum infiltrated for 30 mins. Leaves were incubated at 37 °C overnight followed by clearing with ethanol. Leaves were imaged using a Nikon SMZ1500 stereomicroscope, and photographed with a Qimaging Fast 1394 digital camera. Photographs presented without altering original integrity of images. Similarly, *pTCSv2:GUS* plants were analyzed for GUS expression following *t*Z (10 μM) or DMSO watering. At 60 DAS, excised mature and young leaves were photographed.

*Transcriptomic analysis*

Leaf 5-6 were excised on day of bolting, and flash frozen in liquid nitrogen and stored in -80 °C until grinding. Leaves 5-6 were pooled from three plants (total 6 leaves) for each biological replicate performing a total of three biological replicates. Total RNA was isolated using RNeasy Plant Mini Kits (Qiagen) as per the manufacturer’s instructions and sent off for quality check, library preparation, and sequencing at Novogene Corporation Inc. Following quality check, mRNA was purified from total RNA using poly-T oligo-attached magnetic beads and utilized for cDNA synthesis followed by paired-end 150 bp sequencing on Illumina NovaSeq X-Plus Sequencing Platform. Across all samples, at least 39 million raw reads were generated; which were analyzed with the following bioinformatic pipeline: raw sequence files were checked for quality using FastQC (Andrews, 2010), adapters were trimmed using Trimmomatic (Bolger et al., 2014), and clean reads mapped and counted using HiSAT2 and StringTie (Pertea et al., 2016). TAIR 10 *Arabidopsis thaliana* genome (Araport11 annotation) was utilized as the reference genome. Differential gene expression analysis was performed using DeSEQ2 (Love et al., 2014) after eliminating genes that did not have any mapped reads. Gene ontology (GO) analysis was performed using Shiny GO (Ge et al., 2020) and GO terms with significant enrichment (FDR < 0.05) were retrieved. Raw transcriptome data is available under Bioproject ID: PRJNA1469230.

Resembling the analysis performed in Khanna et al., 2025, WGCNA package was utilized to identify co-expressed gene modules (Langfelder and Horvath, 2008). Briefly, all genes with mapped reads were utilized for DeSEQ2 based variance stabilizing transformation. Using a soft power threshold of 20, adjacency parameters were calculated followed by hierarchical clustering of genes based on pairwise dissimilarity measures. A total of 50 modules were identified and highly similar modules (those with 75% or greater similarity) were merged to yield a total of 45 gene modules. All gene modules were named using a distinct color and visualized using eigengene values. Supp File 5 includes gene module identification, module eigengene expression, and gene module membership values.

*Metabolomic analysis*

Paralleling the transcriptomic analysis, leaf 5-6 across all four genotypes were excised and lyophilized until metabolite extraction was performed across four biological replicates. Metabolomics samples were extracted using a methyl tert-butyl ether/methanol mixture across four biological replicates, following protocols described previously (Dufková et al., 2023, Berková et al., 2026) with minor modifications. In brief, the extracts were processed without phase separation and dried using a vacuum concentrator. The dried residues were subjected to methoximation with methoxyamine hydrochloride (40 mg mL⁻¹ in pyridine), followed by derivatization with N-methyl-N-(trimethylsilyl)trifluoroacetamide (MSTFA). The derivatized samples were analyzed using a Q Exactive GC Orbitrap GC-MS/MS system (Thermo Fisher Scientific) coupled to a Trace 1300 gas chromatograph (Thermo Fisher Scientific) equipped with a Zebron ZB-5MSplus GC column (Phenomenex; 30 m × 0.25 mm × 0.25 μm; 10 m guard column). The temperature program consisted of an initial hold at 70 °C for 5 min, followed by a ramp of 10 °C min−1 to 320 °C, with a final hold at 320 °C for 5 min. Data were processed using Compound Discoverer 3.3 (Thermo Fisher Scientific) and searched against the NIST 2023 library, the GC-Orbitrap Metabolomics Library, and an in-house reference library. Only metabolites meeting strict identification criteria (match score ≥85 and retention index deviation ΔRI <1 %) were included in the final dataset. Prior to statistical analysis, data were normalized to dry weight and log2-transformed. Pairwise differential abundance analyses were conducted using the limma R package (version 3.66.0; Ritchie et al., 2015).

*Integrated transcriptomic-metabolomics analysis*

RNA-sequencing and metabolomic analysis were integrated using the mixOmics package (Rohart et al., 2017). Briefly, transcripts per million (TPM) normalization was performed for all gene counts, amongst which the top 10,000 expressed genes across all samples were utilized for mixOmics integration. Normalized metabolite abundance was used across all metabolites and integrated with TPM – in both cases utilizing three biological replicates for each sample - dropping one extra biological replicate from metabolomics after testing PCA based replicate differences. Initially a multiblock sparse partial least squares discriminant analysis (sPLS-DA) or DIABLO model was utilized following the default package suggestions for number of components and variables required – generating a correlation of 0.96 across genes-metabolites from component 1. Results were visualized using circus and clustered image map (CIM) plots. Further, a global PLS modeling approach was utilized to calculate correlations between the transcriptome and metabolome latent variables. Following this, results from the PLS model were utilized to extract Pearson correlation coefficients of all 10,000 genes with the targeted metabolite of interest (spermidine).

*Statistical Analysis*

All growth and physiological analysis were recorded across at least three independent biological replicates and results presented as average of biological replicates. Unless noted otherwise in figure legend, all statistical significance was performed using ANOVA and Tukey’s HSD. For transcriptome analysis, DESeq2 (Love et al., 2014) was utilized to perform analysis across three biological replicates and Benjamini and Hochberg's approach utilized for adjusting P-values with FDR <0.05 (log2FC = 0) serving as the DEG cutoff. For metabolomic analysis, four biological replicates were utilized, and pairwise differential abundance analyses were conducted using the limma R package (version 3.66.0; Ritchie et al., 2015). Finally, correlations analysis was performed utilizing average of raw values across three biological replicates and reported as Pearson correlation coefficient.
